# CAPTAIN: A multimodal foundation model pretrained on co-assayed single-cell RNA and protein

**DOI:** 10.1101/2025.07.07.663366

**Authors:** Boya Ji, Tingting Hu, Jiawen Wang, Mengmeng Liu, Liwen Xu, Qinhao Zhang, Siyun Zhong, Libo Qiao, Yan Zhang, Shaoliang Peng, Fulong Yu

**Author notes:** These authors contributed equally to this work (Boya Ji, Tingting Hu, Jiawen Wang).

## Abstract

Proteins act as the terminal effectors of cellular function, encoding the phenotypic consequences of genomic and transcriptomic programs. Although transcriptomic profiles serve as accessible proxies, they remain incomplete surrogates for the proteomic landscape that ultimately defines cellular phenotypes. Current single-cell foundation models, however, are trained exclusively on transcriptomes, resulting in biased and partial characterizations of cellular states. To address this limitation, we introduce CAPTAIN, a multimodal foundational model pretrained on over four million single cells with concurrently measured transcriptomes and a curated repertoire of 382 surface proteins across diverse human and mouse tissues. Our results show that CAPTAIN learns unified multimodal representations by modeling cross-modality dependencies and capturing the diversity of cellular states across complex biological contexts. CAPTAIN generalizes robustly across both fine-tuning and zero-shot settings, excelling in core downstream tasks such as protein imputation and expansion, cell type annotation, and batch harmonization. Beyond improved accuracy in multi-omics integration, CAPTAIN uncovers previously inaccessible mechanisms of protein-driven intercellular dynamics, including immune interaction patterns linked to COVID-19 severity. CAPTAIN establishes a new paradigm for multimodal single-cell modeling, laying the foundation for comprehensive cellular understanding and virtual cell construction.

## 1 Introduction

Single-cell sequencing has transformed our ability to define cell states and resolve cellular heterogeneity across biological systems [1–4]. Through large-scale initiatives such as the Human Cell Atlas, hundreds of millions of individual cells have been profiled, yielding expansive transcriptomic landscapes spanning tissues, developmental trajectories, disease conditions, and experimental perturbations [5–8]. This exponential growth of high-dimensional data has catalyzed the development of foundation models [9, 10], which are large-scale self-supervised architectures with self-attention mechanisms trained on massive single-cell corpora [11]. Emerging models such as scGPT [12] and scFoundation [13], alongside novel architectures exploring federated learning for privacy-preserving applications [14], leverage the sequential structure of gene expression data, akin to language models in natural language processing, to distill semantic representations of cell identity that generalize across downstream single-cell tasks [15–19]. While these models have demonstrated strong performance in tasks such as cell type classification, gene expression imputation, and perturbation response modeling, their reliance on transcriptomic data alone often does not fully capture protein-level variation, particularly in immunological and developmental contexts where surface proteins serve as key phenotypic determinants [20, 21]. This unimodal paradigm, although scalable and widely accessible, imposes inherent limitations on our ability to resolve cellular immunophenotypes and capture the full complexity of cell states. Current proteomic profiling methods, with Cellular Indexing of Transcriptomes and Epitopes by Sequencing (CITE-seq) being the most widely adopted, enable the simultaneous measurement of transcriptomes and surface proteins [22, 23]. However, their utility is constrained by the limited and heterogeneous composition of cell surface protein (CSP) panels, which vary substantially between studies and typically encompass only tens to a few hundred targets [24]. Compounded by challenges in cross-modality integration, batch effects from technical and biological sources, and dataset harmonization, these limitations hinder the interpretability of cellular states and the resolution of subtle phenotypic differences [25–27]. These challenges underscore the urgent need for a multimodal modeling framework that integrates transcriptomic and proteomic signals at single-cell resolution, thereby enabling biologically grounded representations that generalize across systems, species, and analytical tasks.

Here, we present CAPTAIN, a single-cell foundation model jointly pre-trained on transcriptomic and proteomic profiles. CAPTAIN (ContrAstive Proteome and Transcriptome pre-trAINing) is designed to bridge the gap between gene expression and protein-level phenotypes by learning cross-modal relationships at single-cell resolution. To enable this, we curated scP&T-4M, a harmonised dataset of over 4.2 million single cells with matched transcriptomes and 382 standardised surface proteins, annotated with a unified nomenclature and functional labels. CAPTAIN is initialized with transcriptomic embeddings learned from tens of millions of single-cell RNA profiles by a large-scale single-cell language model, conferring strong representational capacity. It is further optimized via a cross-attention mechanism to integrate transcriptomic and proteomic features. This provides a rich molecular foundation upon which cross-modal supervision generates protein-informed embeddings that generalise across both unimodal and multimodal contexts. The resulting multimodal joint embeddings support a range of analytical tasks, including clustering, cell type annotation, batch correction and multi-omic integration. Despite being trained on multi-omic data, CAPTAIN accurately predicts surface protein abundance from transcriptomes alone, enabling zero-shot inference across unmeasured targets and extending proteomic interpretability to widely available RNA-only single-cell datasets derived from diverse tissues, conditions, and model systems. We further show that CAPTAIN facilitates biological discovery by reconstructing protein-mediated cell–cell interaction networks and revealing context-dependent surface markers relevant to immune function and disease states.

## 2 Results

### 2.1 Unifying transcriptome and proteome representations with CAPTAIN

CAPTAIN is a single-cell foundation model pre-trained on large-scale human and mouse datasets comprising co-assayed transcriptome and proteome profiles. It adopts a dual-encoder Transformer architecture, in which RNA and protein modalities are processed independently and integrated via a cross-modal attention mechanism to produce a unified representation of cellular state (Fig. 1, Methods and Supplementary Note 4). To enhance transcriptome modeling, the RNA encoder incorporates a gene knowledge module that encodes prior gene–gene relationships and captures high-order regulatory dependencies. Additionally, CAPTAIN leverages transcriptomic embeddings inherited by its RNA encoder from scGPT, a foundational model pretrained on over 50 million single-cell RNA profiles. This initialization yields robust and transferable transcriptomic representations, enabling broad generalization across diverse biological contexts. The protein encoder is trained jointly with the RNA encoder,and the cross-modal attention module captures fine-grained alignment between transcriptional and proteomic features, resulting in a shared embedding space. CAPTAIN simultaneously optimizes unsupervised representation learning for gene expression reconstruction and supervised metric learning for protein abundance prediction, using a multi-objective training framework (Fig. 1a). This design enables the model to learn intricate transcriptome–proteome relationships and to generate modality-agnostic, biologically grounded representations that support robust transfer to diverse downstream tasks (Fig. 1b).

**Fig. 1.**
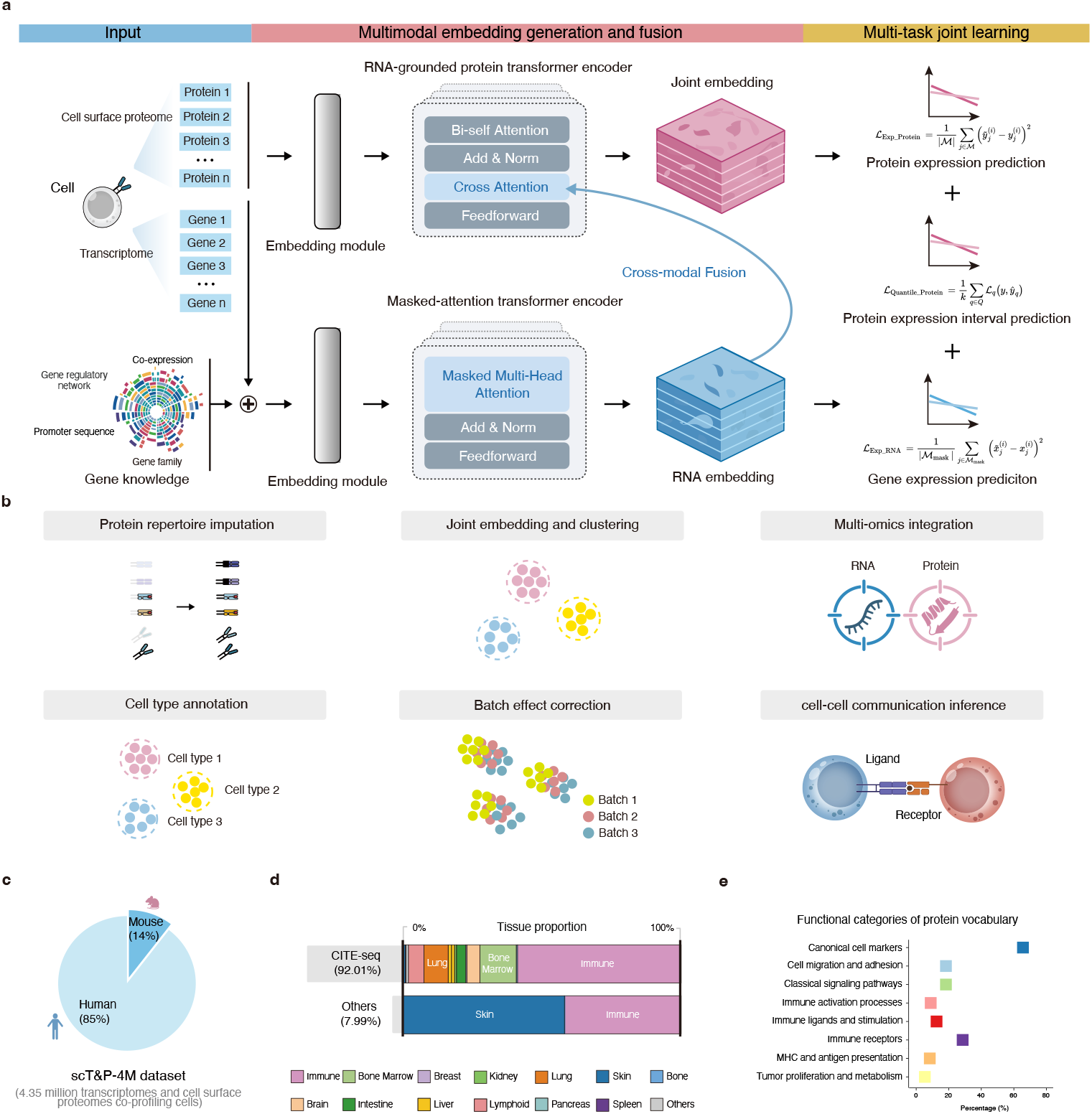
Overview of the CAPTAIN model architecture, training strategy, and pretraining corpus. a.CAPTAIN is pre-trained on large-scale co-assayed single-cell RNA and protein datasets from human and mouse. Gene expression data and prior gene-relevant knowledge are embedded and processed through a masked-attention Transformer encoder to generate contextualized RNA embeddings. Protein expression data is embedded and passed through a protein encoder equipped with both selfattention and RNA-guided cross-attention, producing a joint transcriptome–proteome representation. The model is optimized using a multi-task objective comprising protein abundance regression, interval classification of protein levels, and self-supervised reconstruction of gene expression (Methods). b.The pretrained model enables a wide range of downstream single-cell analysis tasks, including protein imputation and expansion, clustering, multi-omic integration, cell type annotation, batch effect correction, and inference of cell–cell communication. **c.** The curated pretraining corpus, scT&P-4M, comprises over 4.2 million co-profiled single cells with matched transcriptomic and surface proteomic measurements from both human and mouse (Supplementary Table 1). **d.** The dataset spans more than 12 tissues and organ systems, and 11 disease types, supporting the learning of diverse biological representations (Supplementary Table 2 and 3). **e.** After rigorous standardization of nomenclature and functional annotation, 382 distinct surface proteins were used as the proteome vocabulary in CAPTAIN, covering a wide range of molecular functions (Supplementary Table 4-8).

Despite the vast and rapidly expanding repository of single-cell RNA-seq data, corresponding protein-level measurements remain scarce, and inconsistencies in experimental protocols and antibody nomenclature further hinder effective alignment and joint modeling of the two modalities. We therefore curated and harmonized publicly available co-assay datasets (Fig. 1c,d, Supplementary Tables 1-3), resolving naming ambiguities and establishing a unified vocabulary of 382 surface proteins with detailed functional annotation (Fig. 1e, Supplementary Tables 4-8). This effort yielded a comprehensive and standardized corpus of co-assayed single-cell transcriptomic and proteomic profiles, termed scT&P-4M, which provides a robust foundation for crossmodal pretraining. scT&P-4M comprises over 4.2 million single cells from both human and mouse, spanning diverse sequencing platforms, tissue origins, and molecular functions across 252 independent samples. The majority of profiles (92.01%) originate from CITE-seq, with the remainder derived from other compatible co-assay platforms. This resource enables precise cross-modal representation learning and makes possible zeroshot inference of previously unseen proteins, while its broad coverage across tissues, species, and platforms ensures that CAPTAIN learns biologically grounded representations with strong generalization capacity.

A key advantage of CAPTAIN’s contrastively aligned pretraining is its ability to pro-duce generalizable embeddings that can be readily adapted to diverse downstream tasks across both modalities through lightweight fine-tuning. To support this, we developed a flexible fine-tuning pipeline compatible with a broad spectrum of core applications in single-cell analysis, including protein imputation or expansion from transcriptomic inputs, cell type annotation, batch effect correction, multi-omic integration, and cell–cell communication inference (Fig. 1b and Methods). This modular design enables CAPTAIN to serve as a universal backbone for single-cell analytics, supporting scalable deployment across tissues, modalities, and biological contexts.

### 2.2 CAPTAIN enables zero-shot inference of unmeasured surface proteins

Surface proteins are critical for defining cellular phenotype, immune function, and therapeutic targets [22, 28], yet their measurement in single-cell experiments remains both technically demanding and economically costly. Antibody-based modalities such as CITE-seq offer a scalable means to quantify RNA and protein simultaneously, but are constrained by high reagent costs, limited antibody availability, and variable signal-to-noise ratios. As a result, many studies either exclude proteomic profiling altogether or include only a limited subset of proteins. In the scT&P-4M corpus, for instance,protein panel sizes across datasets range from 2 to 210 (mean = 78), more than 4.9-fold smaller than our standardized protein vocabulary. It is therefore critical to assess whether CAPTAIN can accurately infer protein abundance from transcriptomic data, by imputing measured proteins or predicting those that are unmeasured. To this end, we evaluated both the fine-tuned and zero-shot performance of CAPTAIN across four representative CITE-seq datasets spanning different tissues, species, and protein panels (Supplementary Table 9). These included a human peripheral blood mononuclear cell (PBMC) dataset with 168 proteins and 20,729 genes [29]; a human mucosa-associated lymphoid tissue (MALT) dataset with 10 proteins and 33,538 genes (10x Genomics); a human monocyte (MNC) dataset with 10 proteins and 29,929 genes [30]; and a mouse PBMC dataset with 80 proteins and 31,056 genes [31]. For each dataset, we held out a test subset of cells and compared CAPTAIN to three state-of-the-art methods (Seurat [29], sciPENN [30], and TotalVI [32]) using Pearson Correlation Coefficient (PCC) and Root Mean Square Error (RMSE) as evaluation metrics. CAPTAIN consistently achieved the highest protein prediction accuracy across all datasets, as reflected by both PCC and RMSE values (Fig. 2a). In the zero-shot setting, CAPTAIN remained highly competitive, and in the mouse PBMC dataset, it even outperformed all other methods without dataset-specific tuning, reflecting the benefit of its extensive murine pretraining. While Seurat showed slightly higher PCCs in some datasets under the zero-shot setting, CAPTAIN consistently achieved lower RMSE values, indicating more precise predictions with reduced variance. These results suggest that CAPTAIN captures not only the correct directional trends but also provides finer-grained estimations, making it a practical alternative when paired protein data or computational resources are limited.

**Fig. 2.**
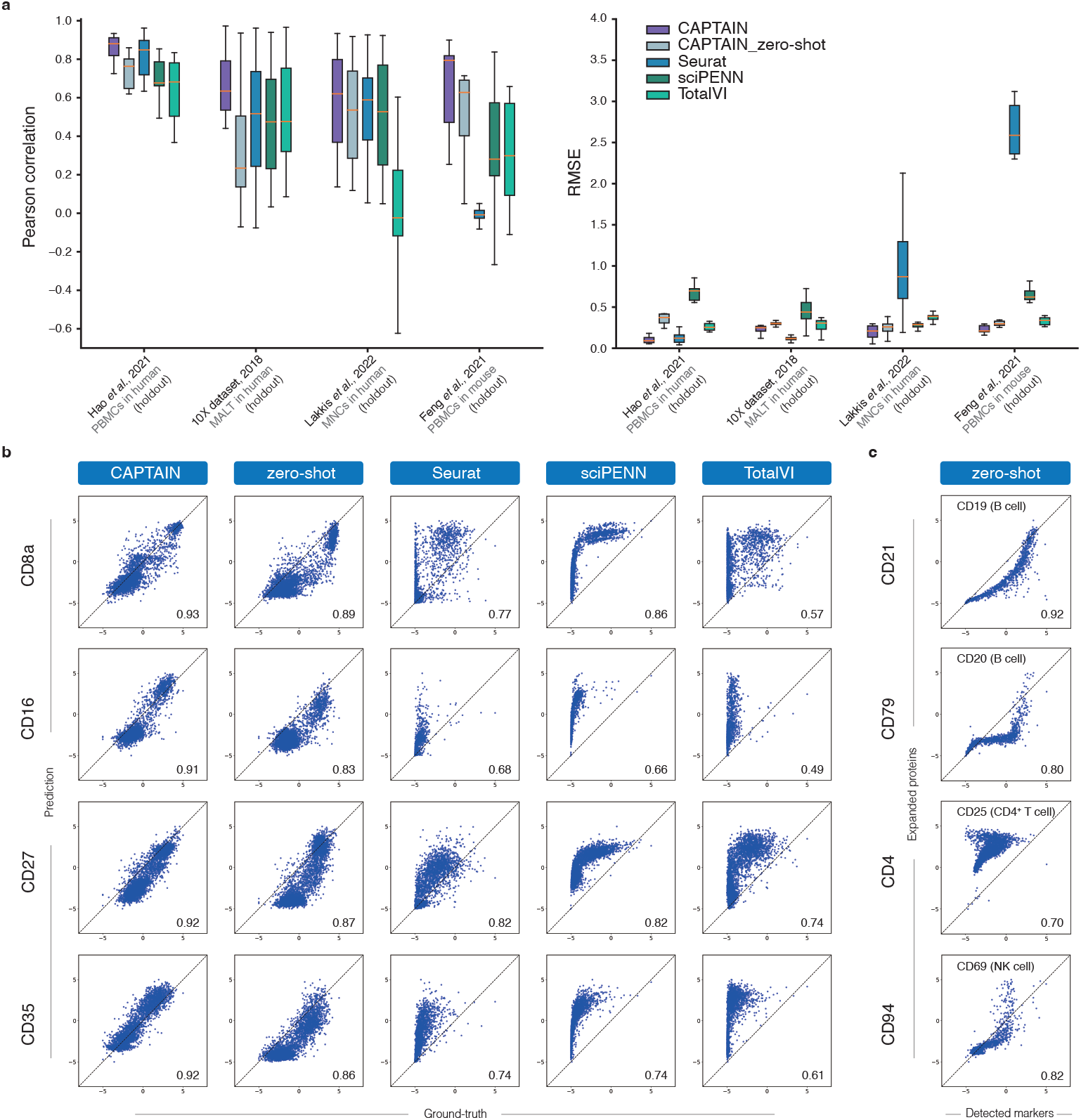
CAPTAIN predicts surface protein expression from transcriptomic inputs in fine-tuned and zero-shot settings. **a.** Prediction performance was evaluated by comparing CAPTAIN with Seurat, sciPENN, and TotalVI across four distinct CITE-seq datasets. Held-out test sets excluded from finetuning were used to evaluate both fine-tuned and zero-shot configurations, with Pearson correlation coefficient (PCC, left panel) and root-mean-square error (RMSE, right panel) as evaluation metrics. Boxes indicate medians, interquartile ranges, and whiskers extend to 1.5 times the interquartile range. **b.** Marker-level prediction accuracy was examined in the human PBMC dataset by comparing predicted and measured expression values for four canonical immune proteins: CD8a, CD16, CD27, and CD35. Predictions from CAPTAIN aligned closely with measured values in both fine-tuned and zero-shot settings, whereas other methods showed greater variance and systematic deviation from the identity line. **c.** Unmeasured proteins that were completely excluded from fine-tuning were predicted by CAPTAIN and evaluated by comparison to the expression of known proxy markers representing the same cell types. CD21 was compared to CD19 and CD79 to CD20 for B cells, CD4 to CD25 for CD4?T cells, and CD94 to CD56 for NK cells.

We further examined CAPTAIN’s predictions for key immune markers in human PBMCs, including CD16, CD8a, CD35, and CD27. The predicted and observed expression values showed strong concordance, with tight clustering along the diagonal and high PCCs in both fine-tuned and zero-shot configurations (Fig. 2b, Supplementary Fig. 1). In contrast, competing methods produced more dispersed predictions with greater error variance. Notably, CAPTAIN was able to accurately predict immune marker proteins that were absent from the fine-tuning data, such as CD21, CD79, CD4, and CD94. The predicted expression patterns of these proteins closely matched the known canonical markers of their respective cell types (e.g., B cells, CD4^+^ T cells, NK cells), despite the absence of any paired protein measurements during training (Fig. 2c). These findings underscore CAPTAIN’s unique zero-shot capability, which enables accurate prediction of surface proteins that are unmeasured or entirely absent in new datasets. This allows substantial expansion of the proteomic feature space from limited-protein or transcriptome-only inputs. Such capability is not found in existing state-of-the-art models and reflects the advantages of CAPTAIN’s biologically grounded architecture and extensive multi-omic pretraining.

### 2.3 Robust cell type annotation across modalities and resolutions

Leveraging its pretraining on large-scale multi-omic data, CAPTAIN enables accurate cell type annotation than models trained solely on scRNA-seq data. To fine-tune CAPTAIN for this task, a neural network classifier was added to process cell embeddings from the protein encoder and output categorical cell type predictions. This composite model was trained end-to-end using cross-entropy loss on a reference dataset with expert-curated annotations and then applied to a separate query partition. CAPTAIN was evaluated across multiple single-cell transcriptomic and multi-omic CITE-seq datasets (Supplementary Table 10). In the human PBMC CITE-seq dataset, UMAP visualizations colored by reference annotations (Fig. 3a) and by CAPTAIN’s predicted labels (Fig. 3b) showed strong concordance, yielding an overall classification accuracy of 96.1%. The corresponding confusion matrix (Fig. 3c) indicated high precision (above 90%) across most cell types. Slight reductions in performance were observed for rare cell populations in the reference set, including CD8^+^ effector memory T (Tem) cells, mucosa-associated invariant T (MAIT) cells, and plasmacytoid dendritic cells (pDCs). To evaluate generalization across modalities and datasets, we expanded the benchmark to five scRNA-seq and four CITE-seq cohorts, including both held-in datasets from the pretraining corpus and held-out datasets not seen during training (Fig. 3d). Five-fold cross-validation was applied to each dataset, and the average macro-F1 score across all cell types was used as the evaluation metric. CAPTAIN consistently outperformed all baseline methods across both modalities, including scGPT, which was evaluated using its published multi-modal extension protocol to ensure a fair comparison with CAPTAIN under equivalent transcriptomic and proteomic input settings. Next, we sought to assess CAPTAIN’s capacity for fine-grained annotation by applying it to a CITE-seq dataset of bone marrow mononuclear cells with literature-defined labels for 13 T cell subtypes [33]. UMAP projections based on reference annotations revealed distinct spatial segregation between CD4 and CD8 subsets, with CD4 subtypes concentrated in the upper right and CD8 subtypes in the lower left (Fig. 3e). Protein expression patterns recapitulated this separation, whereas gene expression alone was insufficient to clearly distinguish the lineages, highlighting the value of surface protein information for resolving closely related cell states (Fig. 3g,h). CAPTAIN accurately predicted the corresponding marker proteins, reinforcing its ability to infer proteomic distinctions from transcriptomic input. It achieved a macro-F1 score of 0.73 across all T cell subtypes, substantially outperforming Seurat (0.61) and scGPT (0.04) (Fig. 3f and Supplementary Fig. 2–4). scGPT misclassified all T cells as CD4^*T*^ naive, while Seurat only partially distinguished CD4^+^ and CD8^+^ groups. These results highlight CAPTAIN’s strength in fine-grained cell type annotation, enabled by its large-scale multi-omic pretraining and incorporation of biological priors, which together yield more discriminative and generalizable representations than unimodal RNA-based models.

**Fig. 3.**
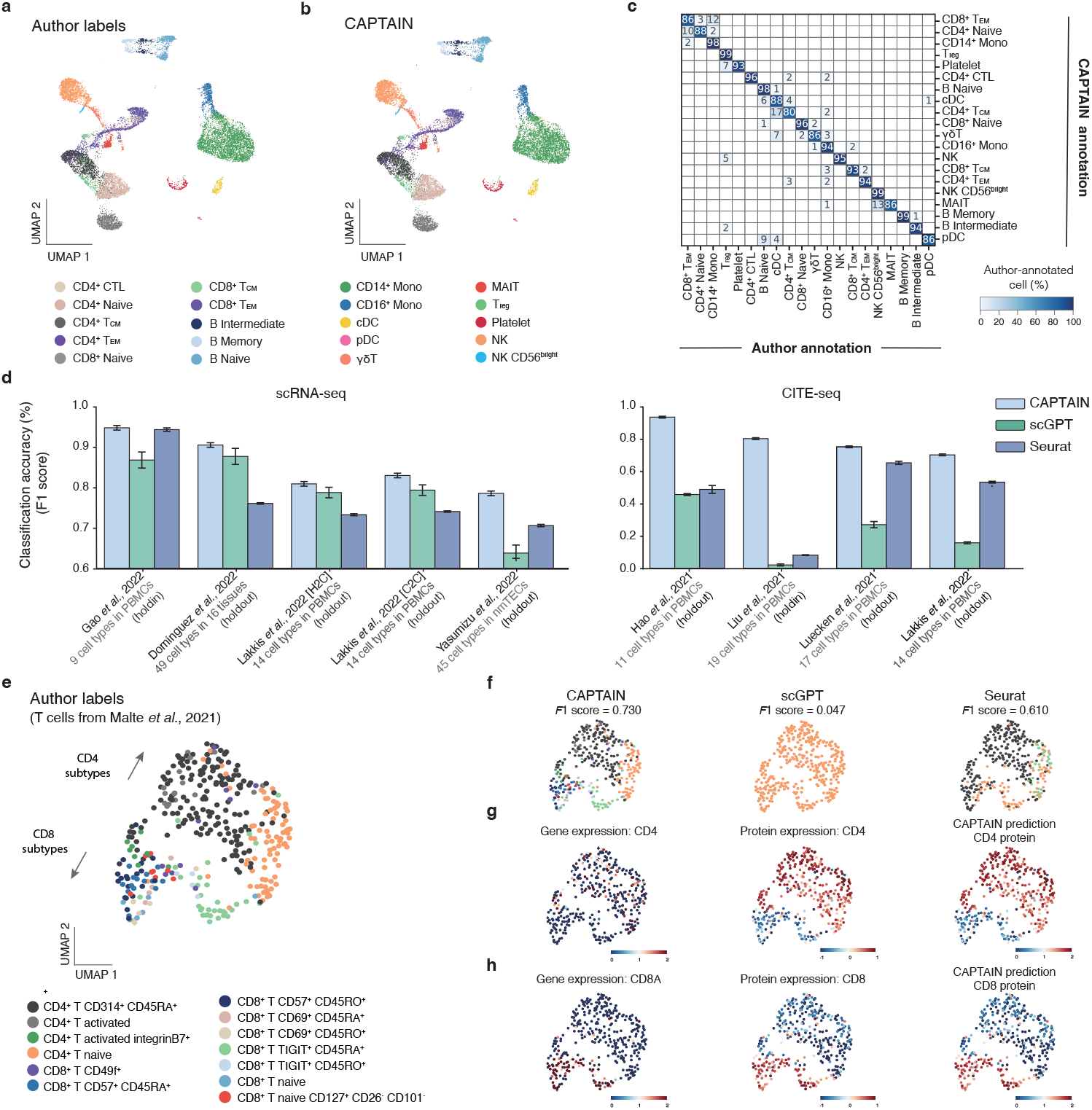
CAPTAIN enables accurate and fine-grained cell type annotation across diverse single-cell datasets. **a.** UMAP visualization of a human PBMC CITE-seq dataset, with cells colored by author-provided reference annotations. **b.** Same UMAP projection as in (a), with cells colored by CAPTAIN-predicted labels, showing strong concordance with the reference. **c.** Confusion matrix comparing CAPTAIN’s predictions to the reference annotations in the PBMC dataset. Each column corresponds to a reference cell type, and each row to a predicted label; values represent the percentage of cells from each true population assigned to each predicted class. **d.** Cell type annotation performance of CAPTAIN, scGPT, and Seurat, evaluated by macro-F1 score across five scRNA-seq datasets (left) and four CITE-seq datasets (right). Datasets include both held-in and held-out samples with respect to CAPTAIN’s pretraining corpus. Bars indicate mean performance across five-fold cross-validation; error bars represent standard deviation. **e.** UMAP visualization of a bone marrow mononuclear cell (BMMC) CITE-seq dataset, colored by fine-grained T cell subtype labels from the original study. **f.** UMAP visualizations of CAPTAIN, scGPT, and Seurat predictions for the same dataset, with macro-F1 scores reported above each panel. **g,h.** Marker-level expression visualizations for CD4 (g) and CD8A (h) in the BMMC dataset. Each panel shows ground-truth gene expression, ground-truth protein expression, and CAPTAIN-predicted protein expression. Protein-level signals offer clearer separation between CD4?and CD8?T cells than gene expression, and CAPTAIN’s predictions closely match true protein distributions.

### 2.4 Biologically meaningful integration across batches, modalities and platforms

Integrating scRNA-seq data remains a persistent challenge, requiring the removal of various spurious variations while preserving true biological differences. To address this, we fine-tuned the RNA encoder of CAPTAIN using a self-supervised masked gene reconstruction objective (Methods), enabling it to learn unified representations across diverse technical and biological conditions.

We first benchmarked CAPTAIN against three widely used integration methods including Seurat [29], Harmony [34], and scGPT [12] on three multi-batch scRNA-seq datasets: PBMCs from 8 healthy donors [29], PBMCs from 2 individuals with SARS-CoV-2 [35], and BMMCs from 13 healthy donors [33] (Supplementary Table 11; Fig. 4a,b). Integration performance was assessed using the AvgBIO score, a composite biological conservation metric combining Normalized Mutual Information (*NMI*_*cell*_), Adjusted Rand Index (*ARI*_*cell*_), and Average Silhouette Width (*ASW*_*cell*_). CAPTAIN achieved the highest AvgBIO score (0.832), substantially outperforming all baselines across datasets (Fig. 4a). On the PBMCs Healthy dataset, CAPTAIN aligned cells across batches while maintaining clear boundaries between all major cell types (Fig. 4b). It also achieved finer resolution of closely related T cell subsets, including CD4^+^ Naive, CD4^+^ Tcm, and CD8^+^ Naive cells, compared to scGPT (Fig. 4b, upper left). Similar results were observed in the SARS-CoV-2 and BMMC datasets, where CAPTAIN again achieved the highest AvgBIO scores, demonstrating robust integration across tissues and disease contexts. In addition to preserving biological structure, CAPTAIN consistently produced competitive batch mixing scores (AvgBATCH), confirming its ability to remove technical artifacts without compromising cellular identity. We next evaluated CAPTAIN’s performance in integrating single-cell multi-omic datasets, which pose greater complexity due to the heterogeneity of molecular layers. Leveraging its multi-modal pretraining, CAPTAIN directly learns joint embeddings from these disparate input types. On a COVID-19 CITE-seq dataset with paired RNA and protein measurements, CAPTAIN outperformed scGPT [12], TotalVI [32], and Scanpy [36], achieving clearer separation of immune subtypes (Fig. 4c; Supplementary Table 12). Similar advantages were observed on a BMMC multi-omic dataset, where CAPTAIN outperformed scGPT [12], Seurat [32], and Harmony [36] in both integration quality and resolution of immunophenotypes (Supplementary Fig. 6).

**Fig. 4.**
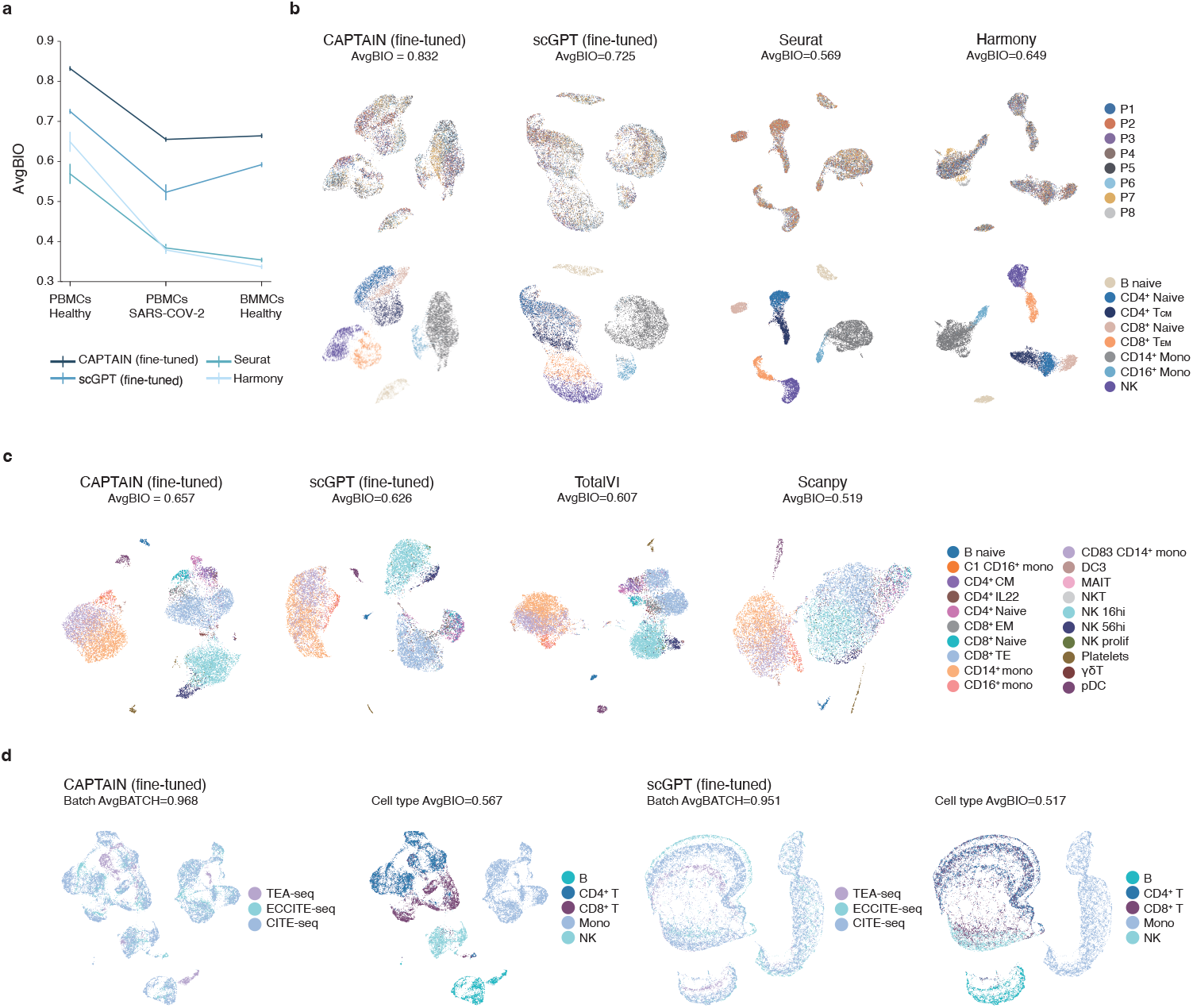
CAPTAIN enables accurate multi-batch and multi-omic single-cell data integration. **a.** Biological conservation scores (AvgBIO) across three multi-batch scRNA-seq datasets: PBMCs from healthy donors, PBMCs from individuals with SARS-CoV-2, and BMMCs from healthy donors. CAPTAIN is benchmarked against scGPT, Seurat, and Harmony. **b.** UMAP visualizations of the PBMCs healthy dataset after integration with each method. Top row shows cells colored by batch (donor), indicating the degree of batch mixing. Bottom row shows cells colored by reference cell type annotations, assessing preservation of biological identity. **c.** UMAP embeddings of the BMMC CITE-seq dataset, generated by CAPTAIN, scGPT, TotalVI, and Scanpy. Cells are colored by fine-grained cell subtype annotations to visualize the quality of integrated cell clustering. **d.** UMAP embeddings from CAPTAIN and scGPT for heterogeneous datasets spanning three multi-omic technologies: CITE-seq, ECCITE-seq, and TEA-seq. For each method, the left panel shows cells colored by technology platform to assess batch alignment, and the right panel shows cell type labels to assess biological structure preservation. AvgBATCH and AvgBIO scores are displayed for each method.

To assess cross-platform integration, we applied CAPTAIN to diverse single-cell datasets generated using three multi-omic technologies: CITE-seq, ECCITE-seq and TEA-seq. These datasets were originally produced independently but were jointly analyzed in a unified embedding space comprising over 60,000 cells (Fig. 4d) [29, 37]. This setting encapsulates the technical breadth of multi-omic platforms, spanning from CITE-seq (RNA and protein) to ECCITE-seq (adding TCR and CRISPR readouts) and TEA-seq (integrating RNA, protein, and chromatin accessibility). Despite substantial assay heterogeneity, CAPTAIN generated coherent clustering across cell types, with clear separation of CD4^+^ and CD8^+^ T cells. Compared to scGPT, CAPTAIN achieved superior performance on both batch correction (AvgBATCH) and biolog-ical conservation (AvgBIO), demonstrating its scalability and generalization across complex multi-platform settings.

### 2.5 CAPTAIN enables protein-informed cell-cell communication inference

Cell–cell communication orchestrates biological processes by transmitting signals through ligand–receptor interactions. These signalling events underlie tissue homeostasis, immune coordination, and therapeutic response. Although surface proteins directly mediate such interactions, most existing computational methods infer intercellular communication solely from transcriptomic data, owing to the widespread availability of scRNA-seq. However, the exclusion of protein information yields an incomplete view of the cellular signalling machinery, particularly the receptor landscape that determines cellular responsiveness [38]. In contrast, protein-informed inference offers a more intrinsic and biologically grounded representation of intercellular communication. To evaluate whether CAPTAIN can overcome the limitations of transcriptome-only models, we developed a hybrid framework that integrates ligand expression from the transcriptome with receptor abundance imputed from the proteome. CAPTAIN first predicts a comprehensive surface protein profile from RNA data using either zero-shot inference or fine-tuning on paired multi-omic datasets. Ligand–receptor interactions are then inferred by combining transcript-derived ligand levels with predicted receptor abundance, and their statistical significance is assessed through permutation testing. This framework enables the reconstruction of signalling networks that more faithfully capture underlying biology (Fig. 5a and Methods).

**Fig. 5.**
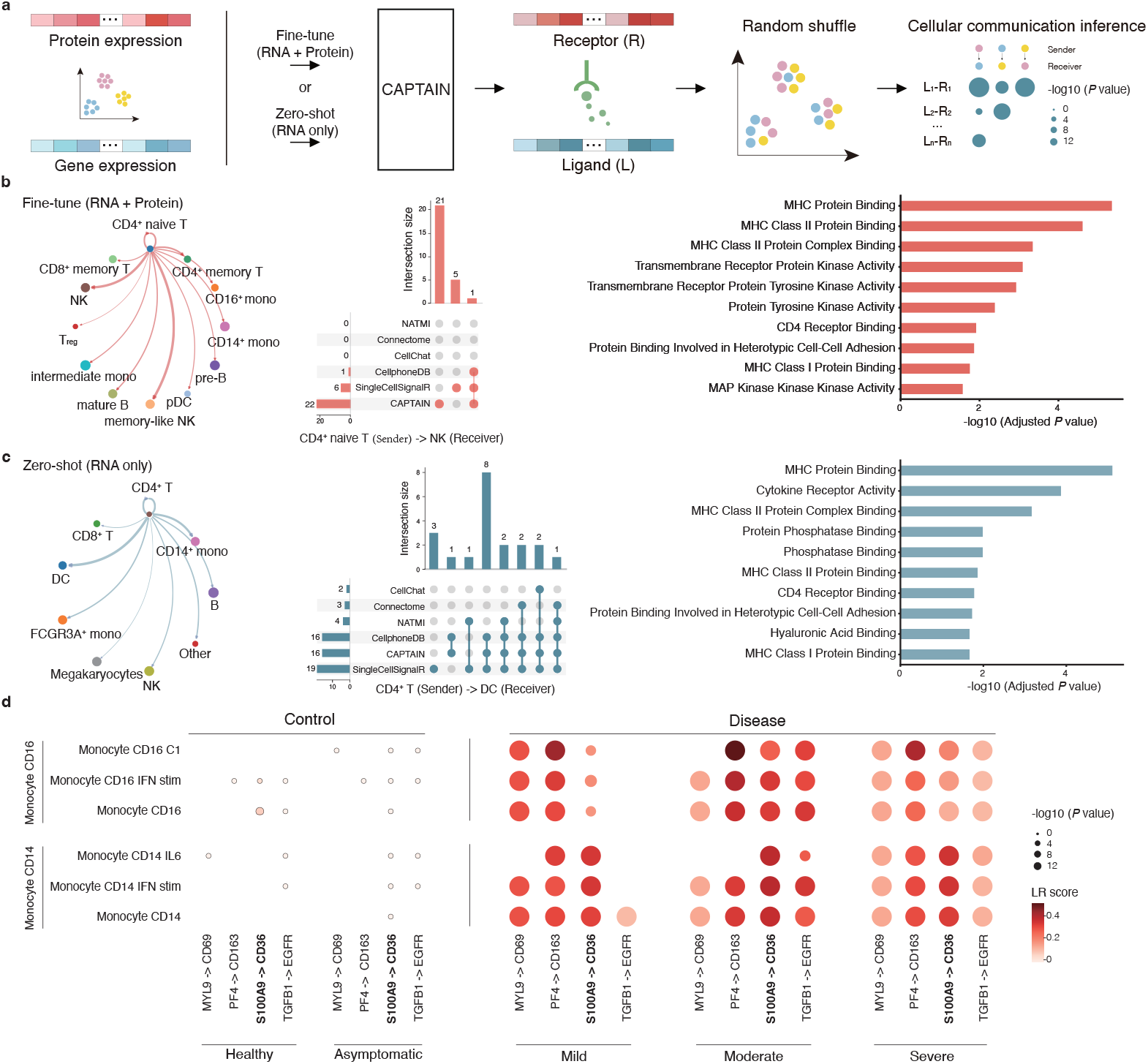
CAPTAIN infers biologically meaningful cell–cell communication from transcriptomic inputs. **a.** Schematic of the CAPTAIN-based communication inference workflow. Ligand expression is extracted from the transcriptome, and receptor abundance is reconstructed using CAPTAIN-predicted protein profiles. Statistical significance of ligand–receptor interactions between cell types is assessed via permutation testing to construct high-confidence signalling networks. **b.** Benchmarking on a CITE-seq PBMC dataset. Left: Circos plot showing inferred signalling from CD4?naive T cells to other cell types. Middle: UpSet plot comparing CD4?naive T - NK cell interactions identified by CAPTAIN and five other methods (NATMI, Connectome, CellChat, CellphoneDB, SingleCellSignalR). Right: GO enrichment analysis of CAPTAIN-inferred interactions highlights immune-relevant molecular functions. **c.** Zero-shot inference on an independent scRNA-seq PBMC dataset without measured surface proteins. Left: Overlap of CD4?T -DC ligand–receptor interactions inferred by CAPTAIN and other methods. Right: GO enrichment confirms functional relevance of CAPTAIN-identified interactions. **d.** Application to a multi-sample COVID-19 dataset. Circos plots show platelet-to-monocyte communication across increasing disease severity (Healthy, Asymptomatic, Mild, Moderate, Severe), highlighting progressive activation of the S100A9–CD36 axis.

We benchmarked CAPTAIN on a CITE-seq PBMC dataset [29] containing paired RNA and surface protein profiles, comparing its performance against five widely used methods: NATMI [39], Connectome [40], CellChat [41], CellphoneDB [42] and SingleCellSignalR [43]. Focusing on CD4^+^ naive T cells, which play a central role in coordinating immune responses [44], we evaluated predicted signalling to NK cells (Fig. 5b, Supplementary Fig. 7). CAPTAIN identified 22 significant ligand–receptor interactions, of which 18 (81.8%) were supported by existing literature. In contrast, the strongest alternative method recovered only 6 interactions, while several others detected none (Supplementary Table 13). GO enrichment analysis of CAPTAIN’s inferred interactions revealed significant overrepresentation of immune-related functions, including MHC protein binding, transmembrane receptor kinase activity, and CD4 receptor binding, underscoring the biological relevance of the predicted communication events.

We further assessed CAPTAIN’s zero-shot capability using another independent scRNA-seq PBMC dataset [35], in which surface proteins were not experimentally measured. CAPTAIN inferred receptor profiles solely from transcriptomic data and used them to predict ligand–receptor interactions between CD4^+^ T cells and dendritic cells (DCs). Of the 16 predicted interactions, 15 were also identified by at least one established method, demonstrating strong concordance with existing tools (Fig. 5c, middle panel). GO enrichment analysis further confirmed the biological relevance of these interactions, revealing significant enrichment for pathways involved in T cell–DC signalling, including cytokine receptor activity and CD4 receptor binding (Fig. 5c, right panel).

To investigate how intercellular communication changes with disease progression, we applied CAPTAIN to a multi-sample COVID-19 dataset [45]. The analysis revealed a progressive increase in platelet-to-monocyte signalling with advancing disease severity, consistent with previous findings [46] (Fig. 5d, Supplementary Fig. 9). Notably, CAPTAIN identified disease-dependent activation of the pro-inflammatory S100A9–CD36 axis, which remained inactive in healthy and asymptomatic individuals but became increasingly prominent in mild to severe cases. In line with prior studies implicating this axis in COVID-19 immunopathology [47], this observation underscores the mechanistic coherence and pathogenic relevance of the ligand–receptor interactions inferred by CAPTAIN.

## 3 Discussion

In this study, we introduce CAPTAIN, a multimodal foundation model trained on a large corpus of co-assayed transcriptomic and proteomic single-cell data. CAPTAIN establishes a unified framework for representing cellular state by integrating transcriptional and surface protein features through a cross-attention Transformer architecture. This design enables it to accurately predict surface protein abundance, perform fine-resolution cell type annotation, integrate diverse single-cell datasets, and infer biologically meaningful cell–cell communication networks. By combining contrastive pretraining with biologically informed modules and multi-objective optimization, CAPTAIN learns high-fidelity, transferable representations that generalize across modalities, tissues, species, and experimental contexts.

A central strength of CAPTAIN lies in its ability to infer surface protein expression directly from transcriptomic inputs. Through comprehensive evaluation across diverse single-cell datasets, irrespective of the availability of paired protein measurements, we demonstrate that CAPTAIN outperforms existing methods in both fine-tuned and zero-shot settings. This capability enables downstream analyses that require multimodal resolution, even in the absence of complete measurements, thereby unlocking tasks that previously relied on explicit multi-omic data. For example, CAPTAIN can reconstruct intercellular communication networks by integrating transcriptomederived ligand expression with imputed receptor abundance, producing biologically coherent signalling maps that capture canonical immune interactions and reveal dynamic changes in disease-associated pathways in COVID-19.

CAPTAIN is pre-trained using a standardized vocabulary of 382 surface proteins curated from over 250 datasets in our scT&P-4M corpus. This unified panel enables consistent modeling across studies and allows CAPTAIN to transcend the sparse and heterogeneous antibody panels typical of current datasets. CAPTAIN substantially expands the effective proteomic feature space available to RNA-only or limited-panel studies, offering a scalable solution to prevailing limitations in single-cell proteomics. In addition, the scT&P-4M corpus provides a robust foundation for cross-modal learning, while CAPTAIN offers a flexible framework for constructing biologically grounded digital cell models. Although the current version focuses on RNA and surface proteins, future extensions incorporating intracellular proteins, chromatin accessibility, or metabolic data will further enhance its versatility. As co-assay technologies continue to evolve, CAPTAIN provides a scalable and adaptable architecture for next-generation integrative single-cell analysis.

Taken together, CAPTAIN offers a generalizable and biologically faithful foundation model for the single-cell community. By combining rigorous multi-omic pretraining with scalable deployment across essential analytical tasks, it lays the groundwork for universal modeling of cellular phenotypes. As multimodal datasets grow in depth, resolution, and diversity, we anticipate that CAPTAIN will enable deeper insights into cell identity, state transitions, and intercellular dynamics, advancing the goal of constructing truly comprehensive virtual cells [48, 49].

## 4 Methods

### 4.1 Pre-training data collection and preprocessing

To generate a high-quality pre-training dataset for the CAPTAIN foundation model, we systematically collated and processed publicly available co-assayed single-cell RNA and protein data to construct a scT&P-4M database (Supplementary Note 1). This process comprised three main stages: data acquisition and selection; gene and cell surface protein (CSP) symbol unification; and cell-level quality control and integration. This harmonisation procedure was essential to achieve structural consistency and comparability between the RNA and protein modalities across diverse projects and platforms.

#### Data acquisition and selection

We performed a comprehensive screen of recent studies, using public repositories including the Gene Expression Omnibus (GEO) [50], ArrayExpress [51] and the official 10x Genomics website. Inclusion criteria were as follows: (i) the dataset had to contain both RNA and CSP modalities; (ii) raw count matrices had to be available in a standard format (for example, CSV, H5AD, RDS, or FASTA files subsequently processed using CellRanger); (iii) complete CSP annotation was required, with a preference for datasets containing cell-type annotations; and (iv) sample batch and sequencing platform information had to be documented. This curation process yielded a total of 252 samples, comprising 4,531,996 single cells and 382 distinct CSP features. The final dataset thus contained paired RNA and protein expression profiles from each cell for pre-training.

#### Gene and CSP symbol unification

For the RNA modality, gene symbols in each dataset’s raw count matrix were mapped to a reference vocabulary derived from the HUGO Gene Nomenclature Committee (HGNC) database [52]. This vocabulary, comprising 19,264 protein-coding and mitochondrial genes, was selected to match the feature space of the pre-trained scGPT model [12]. To maintain a consistent feature space across all samples, any gene from this reference set not present in a particular sample was included with a count of zero. To resolve the considerable heterogeneity in CSP naming conventions across studies, we implemented a comprehensive normalisation pipeline for all CSP features (Supplementary Note 2). Common discrepancies included inconsistent prefixes or suffixes derived from fluorophore or batch information (for example, ‘CD45RA-BV421’ versus ‘CD45RA’) and variable capitalisation (for example, ‘CD3’ versus ‘Cd3’). To address these inconsistencies, we developed and applied custom scripts using regular expressions to parse and standardise the CSP labels. Our pipeline specifically removed substrings following hyphens (e.g., ‘CD19-BV786’ became ‘CD19’), stripped product codes, converted all names to uppercase and manually mapped synonymous markers (e.g., ICOS to CD278). This procedure yielded 382 high-quality, standardised CSP features, establishing a robust foundation for subsequent feature alignment and multimodal fusion.

#### Cell-level quality control and integration

For each dataset, we first ensured that both RNA and CSP data were available for the same cell barcodes and retained 15 only these paired cells for subsequent analysis. To filter contaminated empty droplets, extremely low-quality cells and damaged cells, we kept cells with over 200 genes expressed (that is, expression vector with nonzero value count *>*200) for pretraining by using the Scanpy [36] packages. Following this quality control step, the processing pipelines for the two modalities diverged. The RNA expression matrix for each cell was normalised to a library size of 10,000 and subsequently log-transformed. The corresponding CSP count matrix was separately normalised and then scaled to have zero mean and unit variance, aligning with established common practices for processing CSP counts from CITE-seq experiments [29, 30].

Finally, the processed data from both modalities were consolidated into a single data object. This object contained key features for each cell, including expression matrices, UMAP coordinates and associated metadata. Where available, cell-type annotations were integrated using a left join on cell identifiers. The final data were stored in the H5MU format, which served as the standard input for model pre-training. The purpose of this pipeline was to generate a standardised and harmonised dataset suitable for the cross-modal pre-training tasks described in the following sections.

### 4.2 CAPTAIN model architecture

We leveraged the Contrastive Language-Image Pre-training (CLIP) model [53] as the backbone model of CAPTAIN. The transcriptomic module receives four primary inputs: a priori gene knowledge, species tokens, gene tokens, and their corresponding expression values. In parallel, the proteomic module processes cell surface protein tokens. These distinct inputs are transformed by dedicated embedding layers before being encoded by separate transformer architectures. Within the transcriptomic pathway, a masked-attention transformer, inspired by scGPT [12], captures critical gene-gene and gene-cell relationships. The resultant RNA embeddings then guide a cross-attention mechanism within the proteomic encoder to inform the prediction of protein abundances from their corresponding cell surface protein tokens. This RNA-informed process culminates in a unified, multi-omic joint embedding that supports a range of downstream analytical applications.

#### 4.2.1 Prior biological knowledge

To enrich our pre-training model, we integrated four modalities of prior biological knowledge for human and mouse genes, sourced from the GeneCompass framework, whose embeddings have been demonstrated to enhance the learning of cross-species homology [54]. Each modality was encoded into a common 512-dimensional embedding space. The specific embeddings utilised were:

##### Gene Regulatory Network (GRN) embeddings (*emb*_*grn*_)

These were derived from 84 mouse and 76 human GRNs that had been constructed using paired gene expression and chromatin accessibility data from the Encyclopedia of DNA Elements (ENCODE) [55]. The embeddings represent regulatory gene pairs from these networks, generated via the gene2vec algorithm [56].

##### Promoter embeddings (*emb*_*pro*_)

These were generated by fine-tuning a pre-trained 16 DNABert model [57] with promoter sequences spanning 2,500 bp around the transcription start site (500 bp upstream and 2,000 bp downstream).

##### Gene family embeddings (*emb*_*fam*_)

Human gene family data were sourced from the HUGO Gene Nomenclature Committee (HGNC) [52]. Mouse gene families were inferred via homology using BioMart (Ensembl) [58], resulting in 1,645 and 1,539 families for human and mouse, respectively. Within each family, pairs of genes were defined and subsequently embedded using the gene2vec algorithm [56].

##### Co-expression embeddings (*emb*_*coe*_)

These were established from Pearson Correlation Coefficients (PCCs) calculated using uniformly sampled single-cell gene expression data. The final embeddings represent gene pairs with a PCC*>*0.8, generated via the gene2vec algorithm [56].

#### 4.2.2 RNA embedding module

Our RNA embedding module adapts the scGPT architecture [12], primarily to leverage its publicly available weights pre-trained on over 33 million cells. This initialisation provides a robust baseline for gene representation. In contrast to the original framework, our module further incorporates specific prior biological knowledge to generate more biologically informed gene embeddings. The module input is derived from the raw cell-by-gene count matrix, denoted as *X*∈ ℝ^*N* ×*G*^. In this matrix, rows correspond to cells (*i* = 1, *…, N*) and columns to genes (*j* = 1, *…, G*). For each cell *i*, its input comprises the following components.

##### Gene Tokens 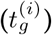

In this framework, each gene is treated as a fundamental token of biological information, analogous to a word in natural language processing. A single, unified vocabulary is constructed to enable the harmonisation of data from diverse sources, such as studies employing different sequencing technologies or preprocessing pipelines. This is achieved by taking the union of all unique genes across the relevant datasets, after which each gene *g*_*i*_ in the global vocabulary is assigned a unique integer identifier, *id*(*g*_*i*_). A special < *pad* > token is also introduced to ensure all input sequences have a fixed length through padding. Following this tokenization, each cell *i* is represented as a sequence of *M* integer identifiers in a vector 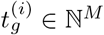:

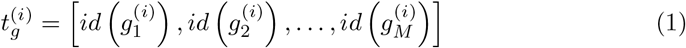

where *M* is a predefined hyperparameter for the maximum input length.

##### Species Tokens 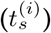

To explicitly inform the model of the species of origin for each cell, a species-specific token is incorporated into the input representation. This enables the model to learn and leverage species-specific biological patterns during pre-training. The species identity is encoded using a simple integer mapping, where human is assigned an identifier of 0 and mouse an identifier of 1. This identifier is then broadcast across the maximum sequence length, *M*, to create a species token vector, 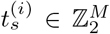, for each cell *i*. This vector is defined as 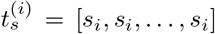, where the species identifier *s*_*i*_ is determined by the origin of cell *i*:

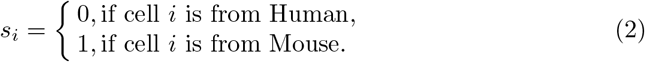

##### Gene expression values (*x*^*i*^)

To address the variability in expression magnitudes arising from factors like sequencing depth and gene sparsity, the value binning technique is adopted [12]. This method normalises expression counts onto a relative, cell-specific scale. The procedure is applied on a per-cell basis. To speed up the pretraining, we restrict the input to only genes with non-zero expression for each input cell. For each cell *i*, its non-zero expression values *X*_*i,j*_ (*X*_*i,j*_ *>* 0) are sorted and partitioned into *B* quantiles of equal size. This process defines *B* consecutive, cell-specific intervals [*b*_*k*_, *b*_*k*+1_], where *k ∈* 1, 2, *…, B*. The raw expression count *X*_*i,j*_ is then mapped to a binned integer value 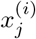 according to the rule:

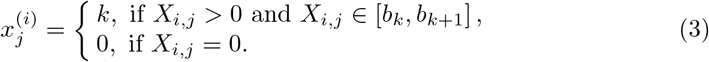

This ensures the resulting values have a consistent semantic meaning; for instance, a binned value of *B* consistently represents the highest expression tier within that specific cell. For certain fine-tuning applications, this binning step is preceded by log1p transformation and highly variable gene (HVG) selection. The final vector of binned values for cell *i* is denoted as 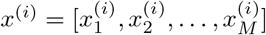.

##### Input Embeddings

The gene tokens and species tokens, *emb*_*g*_ and *emb*_*s*_, are embed-ded using conventional pytorch embedding layers (torch.nn.Embedding) to facilitate the mapping of each token to a fixed-length embedding vector of dimension *D*. Concurrently, binned expression values are processed by a multi-layer perceptron (*emb*_*b*_), which captures the ordinal nature of gene expression values. The final embedding for cell *i, h*^(*i*)^ *∈ ℝ*^*M ×D*^, is the element-wise sum of these component vectors:

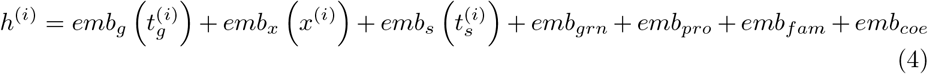

#### 4.2.3 Protein embedding module

This module encodes a panel of 382 curated cell surface protein tokens. Each token is assigned a unique integer identifier, *id*(*p*_*j*_), to form a vocabulary. A special classification token, *< cls >*, is included to aggregate the protein-level embeddings into a global cell representation. For a given cell, *i*, the sequence of its protein tokens is represented as a vector,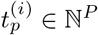 :

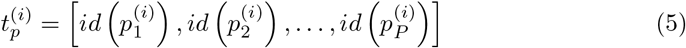

where *P* =382. These protein tokens are embedded using conventional pytorch embedding layers, 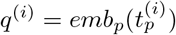, which project each discrete identifier into a fixed-length embedding vector of dimension *D*.

#### 4.2.4 RNA encoder module

To address the non-sequential nature of single-cell data and effectively leverage the pre-trained parameters derived from scGPT [12], we employed a masked-attention transformer architecture to encode the complete input RNA embeddings, denoted as *h*^(*i*)^ (Equation 4). The transformer’s self-attention mechanism operates on sequences of *M* embedding vectors, allowing it to capture putative gene-gene interactions. The output of the final *n*_*th*_ transformer layer, the resulting RNA representation, *h*_*n*_^(*i*)^ = *Transformer*(*h*^(*i*)^)*∈* ℝ^*M* ×*D*^, is then fused with corresponding protein embeddings via a cross-attention module to generate a robust multi-modal embedding. To ensure computational efficiency when processing large gene panels (where the input dimension *M* can be in the tens of thousands), the model employs an accelerated selfattention mechanism, FlashAttention. This memory-efficient implementation allows for the feasible and effective application of self-attention to extensive genomic data. Furthermore, standard autoregressive models are ill-suited for gene expression data, which, unlike natural language, is non-sequential and lacks an inherent order for prediction. To address this challenge, we utilized the specialized attention masking strategy to enable autoregressive-style generation. The approach is to predict expression values for a subset of ‘unknown’ genes based on a set of ‘known’ genes. This is governed by a custom attention mask, *A*_*mask*_, which modifies the standard attention equation:

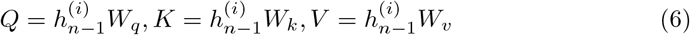

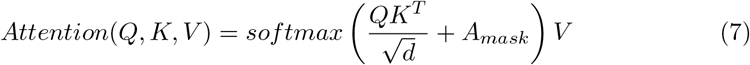

where *Q, K, V* ∈ ℝ^*M* ×*d*^ represent the query, key and value vectors. *W*_*q*_, *W*_*k*_, *W*_*v*_ ∈ ℝ^*D*×*d*^ are the learnable weight matrices. Specifically, the mask prohibits a given query from attending to any ‘unknown’ genes other than itself by setting the corresponding weights in the attention matrix to *−∞*. This is defined for each element *a*_*i,j*_ in *A*_*mask*_

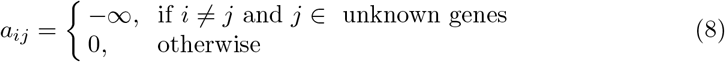

Generation is performed iteratively. In each step, the model predicts expression values, and the most confidently predicted genes are added to the ‘known’ set for the next iteration. This workflow effectively mimics causal prediction for non-sequential data and unifies generation from both gene-level and cell-level prompts.

#### 4.2.5 Protein encoder module

The protein encoder module uses a canonical Transformer-encoder architecture, augmented by a dedicated cross-attention block to fuse RNA and protein information into a unified representation. The parameters of this module are initialised at random.

Given an initial protein embedding matrix, 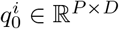, where *P* is the number of protein tokens and *D* is the embedding dimension, we first apply a standard self-attention and feed-forward layer. For *n*_*th*_ layer, the output of the self-attention mechanism, *Self Attention*_*n*_, is computed from the previous layer’s output,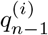:

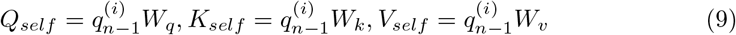

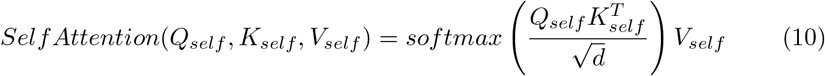

The output of the attention head is then passed through a residual connection, layer normalization, and a standard position-wise feed-forward network. After such block, the final output is 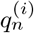, which represents an embedding that encodes intra-protein relationships.

After that, to integrate RNA context, we introduce a cross-attention layer that uses the precomputed RNA embedding *h*^(*i*)^ ∈ ℝ^*M* ×*D*^ as both queries and keys, while treating the protein-encoded features *q*^(*i*)^ ∈ ℝ^*P* ×*D*^as values. Concretely, we compute:

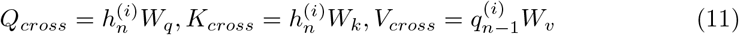

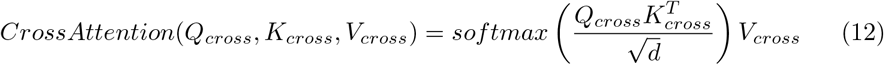

This cross-attention block produces a matrix of size ℝ^*P ×d*^, which is then projected back to ℝ^*P ×D*^, merged with 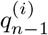 via a residual connection, and normalized. The result is a joint RNA-protein embedding *q*^(*i*)^ ℝ^*P* ×*D*^that simultaneously captures transcriptomic and proteomic features, ready for downstream tasks.

#### 4.2.6 Cell-level representation

A unified, cell-level representation, 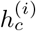, is generated by learning to aggregate features across both the transcriptomic and proteomic modalities. To facilitate this learned aggregation, we introduce a special classification token, < *cls* >, which is prepended to the input sequence of protein tokens. This token, along with the protein features, passes through the protein-specific self-attention encoder and is subsequently fused with the RNA embedding via the cross-attention module. The final hidden state from the fusion block corresponding to this < *cls* > token is then taken as the definitive multi-modal cell representation. This single vector, 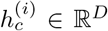, thereby encapsulates the salient, cross-modally informed features of the cell for all downstream predictive tasks.

### 4.3 Learning objective for pretraining

The model is pretrained using a composite objective function designed to learn multimodal representations from single-cell data. This objective combines three distinct components: (1) a masked gene expression prediction task within the RNA modality; (2) a cross-modal prediction of protein expression from RNA features; and (3) uncertainty quantification for the protein predictions via interval estimation. Collectively, these components enable the model to learn both intra-modal gene regulatory patterns and inter-modal transcript-protein relationships.

#### Masked gene expression prediction

The masked gene expression (MGE) prediction objective is utilized to learn gene-gene interdependencies. This task requires the model to infer the expression levels of a randomly masked subset of genes based on the expression of the remaining, unmasked genes within a given cell. During training, the expression values for a random fraction of genes in each cell, *i*, are masked. The model’s RNA encoder then processes the available transcriptomic information to produce contextualised embeddings. Subsequently, a multi-layer perceptron (MLP) head is used to predict the expression value,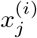, for each masked gene, *j*, from its corresponding hidden state representation,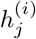. The MGE objective is to minimise the mean squared error (MSE) between the predicted and actual expression values. This loss function, denoted *L*_*MGE*_, is defined as:

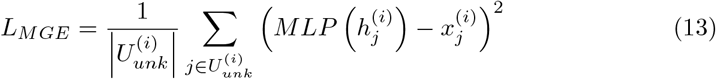

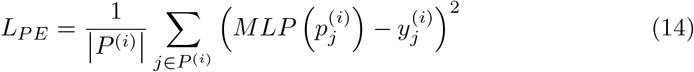

Here, *U*_*unk*_ is the set of indices for the masked genes in cell *i*; 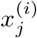 is the groundtruth expression for the *j − th* masked gene; and 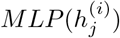 is the value predicted by the MLP. The denominator 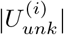 is the total number of masked genes. Optimising this objective enables the model to learn the biological relationships and dependencies within the gene expression data.

#### Protein expression prediction

To align transcriptomic and proteomic representations within a shared embedding space, the model is trained with a cross-modal objective. This objective requires the model to predict the abundance of cell-surface proteins using only the cell’s gene expression features as input. For each cell, *i*, the model predicts the expression level, 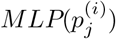, for every protein, *j*, that was measured in that cell’s panel, denoted *P*_*i*_. The learning objective is to minimise the MSE between these predictions and the ground-truth protein expression levels, 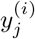, This cross-modal prediction loss, *L*_*P E*_, is therefore defined as:

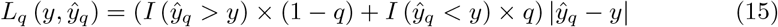

Here, |*P* ^(*i*)^ |is the number of proteins measured in cell *i*. By calculating the loss exclusively over the set of available proteins, this objective robustly handles the incomplete protein panels common in CITE-seq data. Optimising this objective compels the model to learn a joint representation where transcriptomic states are predictive of proteomic outcomes.

#### Protein expression interval prediction

We are interested not only in predicting protein expression but also in quantifying the uncertainty of our prediction using interval estimation. To that end, we will want to estimate not only the mean expected protein expression given the RNA expression profile of the cell but also a vector of quantiles that can be used to construct prediction intervals. Let {*Q* = *q*_1_, *q*_2_, *…, q*_*k*_} be the set of quantiles we wish to estimate. Then we want to estimate the protein expression *y* and the predicted protein expression quantile ŷ_*q*_ for *q ∈ Q* such that we minimize the following objective:

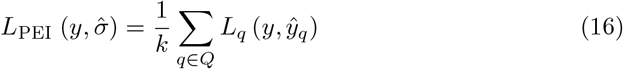

where *I* represents the indicator function. The role of the indicator function here is to apply an asymmetric penalty. It selectively uses either the weight *q* or 1 *− q* depending on whether the prediction is an overestimation or an underestimation, which is the core principle of quantile regression.

### 4.4 Fine-tuning on downstream tasks

#### 4.4.1 Single-cell protein imputation and expansion

For the single-cell protein imputation and expansion tasks, datasets from various tissues were randomly partitioned into training (90%) and test (10%) subsets. We retained the intersection of gene markers common to both our pre-trained foundation model and the reference set. Prior to fine-tuning, all data preprocessing was performed using the Scanpy package. Gene expression values were normalised, log-transformed, and subsequently binned. Paired protein expression values were separately normalised and scaled. Only gene markers with non-zero expression values were utilised for training.

We demonstrate the utility of the pre-trained CAPTAIN model for single-cell protein prediction through two primary downstream applications. In the first application, protein imputation, the model was fine-tuned on paired single-cell transcriptomic and proteomic data. The objective was to impute the expression levels of specific proteins that were missing from our reference CSP panel. In the second application, protein expansion, we employed the model for zero-shot prediction. Here, the model predicted the expression of all proteins in our reference panel using only the cellular transcriptome as input, without requiring paired protein measurements for the test cells. For both applications, the model was initialised with our pre-trained weights. The finetuning process mirrored the parameter settings of the pre-training phase: we used a learning rate of 1 *×* 10^−4^ with a decay to 90% of its value after each epoch, and a batch size of 20. The fine-tuning objective function was identical to that used during the model’s pre-training.

We benchmarked CAPTAIN against three contemporary methods for single-cell protein prediction: sciPENN [30], Seurat (v4) [29], and TotalVI [32]. For Seurat and TotalVI, we followed their standard, published prediction workflows for the comparison. To evaluate performance, we calculated two key metrics for each protein by comparing its measured and predicted expression values across all cells: the Pearson correlation coefficient, to assess the linear relationship, and the Root Mean Square Error (RMSE), to quantify the magnitude of prediction error.

#### 4.4.2 Cell type annotation

For the cell type annotation task, datasets from diverse tissues with manually curated cell type labels were randomly partitioned into training (90%) and test (10%) subsets. This procedure was repeated using multiple random seeds to ensure reproducibility. The model was subsequently fine-tuned on the training data, and its annotation performance was evaluated on the held-out test data. The input features were restricted to the intersection of gene markers common to our pre-trained foundation model and the reference dataset. Prior to fine-tuning, we used Scanpy to normalise, log-transform, and bin gene expression values, and to separately normalise and scale paired protein expression values. All gene markers, including those with zero expression, were retained for the training process.

Our model is designed to perform cell type annotation on two distinct data modalities: single-cell transcriptomes alone (scRNA-seq) and paired single-cell transcriptomic and proteomic data (e.g., CITE-seq). The core architecture remains consistent, with the primary distinction being the placement of the output classifier. For scRNA-seq data, a multilayer perceptron (MLP) classifier is appended to the final layer of the RNA encoder module. For paired multi-omic data, this MLP classifier is instead connected to the final layer of the CSP encoder module. To begin the fine-tuning process, the model was initialised with weights from the pre-trained foundation model. The only exception was the output MLP classifier, whose weights were randomly initialised. Fine-tuning was initiated with a learning rate of 1 *×* 10^−4^ and a batch size of 26. The learning rate was scheduled to decay after each epoch to 90% of its value from the previous epoch. The training objective was to minimise the cross-entropy loss between the predicted cell type probabilities, derived from the learned cellular representations, and the ground-truth cell type labels.

We benchmarked the performance of our model against two contemporary cell type annotation methods: scGPT [12] and Seurat (v4) [29]. The comparison was conducted on multiple datasets using a five-fold cross-validation scheme. Performance was assessed using four standard classification metrics: accuracy, precision, recall, and the macro F1-score. Accuracy, precision, and recall were calculated globally to provide a measure of overall performance. In contrast, the macro F1-score was calculated as the unweighted mean of the F1-scores for each cell type class, thereby giving greater importance to the correct classification of rare cell populations (Supplementary Note 3). For the scGPT comparison, we used the standard scGPT model for scRNA-seq-only datasets and the specific scGPT variant designed for multi-omic integration when analysing CITE-seq data. Both models were trained and used for prediction with default parameters. For the Seurat comparison, we employed distinct workflows depending on the data modality. For scRNA-seq data, we utilised the TransferData function. This method identifies anchors between the reference (training) and query (test) datasets based on principal component analysis (PCA) of the training data, subsequently transferring the known cell type labels to the test set. For multi-omic datasets, we first applied its weighted nearest neighbour (WNN) analysis to learn a joint representation of the RNA and protein modalities. Following this integration, we applied the TransferData workflow on the resulting integrated space to transfer labels from the training to the test set. In both cases, the transferred labels were taken as the predicted cell types for the test data.

#### 4.4.3 Batch correction

When combining datasets from different sequencing batches or technologies, batch effects can be a major issue for downstream analysis, such as cell type clustering. Our goal was therefore to correct for this technical noise in both single-modality and multimodality datasets, while preserving the true biological variance. All datasets were split into training (90%) and test (10%) subsets. The preprocessing method for both singlecell transcriptomic and proteomic data was the same as in the previously described tasks. For model input, we also used 3,000 HVGs with non-zero expression for the RNA encoder module and all available proteins for the protein encoder module.

To correct for batch effects, we fine-tuned the model after loading all pre-trained weights. The cell representation was taken from different sources depending on the input data. For single-modality transcriptomic data, the embedding of the < *cls* > token from the RNA encoder module was used. For multi-omic datasets, this representation was instead taken from the < *cls* > token of the Protein encoder module. We used a multi-task fine-tuning strategy to ensure effective batch integration. We continued to use the GEP and GEPC objectives in the multi-omic integration task. In addition, we added three more objectives to improve performance: Elastic Cell Similarity (ECS), to improve cellular contrastive learning, as well as DAR and Domain-Specific Batch Normalization (DSBN) for direct batch correction [12]. Following the default parameters for these objectives, the masking ratio for GEP and GEPC was set to 0.4, and the *β* parameter in ECS was set to 0.6. Fine-tuning was started with a learning rate of 1 *×* 10^−3^, and the learning rate decayed to 90% of its value after each epoch. The model was trained for a fixed 15 epochs using a batch size of 16. We compared the batch correction performance of CAPTAIN against three other common methods: scGPT [12], Seurat [59], and Harmony [34]. To ensure a thorough evaluation, performance was measured using the same set of biological conservation and batch-correction metrics as detailed in the multi-omic integration task.

#### 4.4.4 Multi-omics integration

For the multi-omic integration task, our model focused on integrating single-cell transcriptomics and proteomics. To evaluate its performance robustly, we used both single-batch and multi-batch multi-omic datasets. Each dataset was also partitioned using a 9:1 ratio. The preprocessing pipelines for the single-cell transcriptomic and proteomic data were identical to those described in the preceding tasks. For the RNA encoder module, the 3,000 most highly variable genes (HVGs) with non-zero expression were selected for training.

We initialised the model with all parameters loaded from the pre-trained foundation model. The final multi-omic representation for each cell was derived from the embedding of the special < *cls* > token within the CSP encoder module. Notably, to facilitate both batch effect removal and effective integration, we adopted several fine-tuning objectives for the RNA encoder module as proposed by scGPT. These included Gene Expression Prediction (GEP), Gene Expression Prediction for Cell modelling (GEPC), and Domain Adaptation via Reverse back-propagation (DAR) [12]. The fine-tuning process was initiated with a learning rate of 1 *×* 10^−4^, which was scheduled to decay to 95% of its value after each epoch. The weight for the DAR objective was set to the default value of 1.0. The model was trained for a fixed 25 epochs with a batch size of 16.

We benchmarked CAPTAIN against recent multi-omic integration methods, including scGPT [12], Scanpy (MOFA) [60], and TotalVI [32], on both single-batch and multibatch datasets. For a fair comparison, all four methods were trained using an identical input feature set, comprising 3,000 HVGs from the transcriptomic data and all available proteins from the proteomic data. The three competing methods were run using their publicly available, default workflows. The quality of the resulting cell embeddings was assessed using two distinct sets of metrics. First, to evaluate the preservation of biological identity, we used four biological conservation metrics [61]: Normalized Mutual Information (*NMI*_*cell*_), Adjusted Rand Index (*ARI*_*cell*_), and Average Silhouette Width (*ASW*_*cell*_). These scores measure the consistency between the derived cell type clusters and the ground-truth labels. We also computed *AvgBIO*, the average of these three metrics, for an aggregate score. Second, given that our datasets included multiple batches, we further evaluated the integration performance using three batch-correction metrics: *ASW*_*batch*_ and Graph Connectivity (*GraphConn*). These metrics assess the degree of mixing between different omic batches. To summarise the batch-mixing performance, we calculated *AvgBATCH*, representing the average of *ASW*_*batch*_ and *GraphConn* (Supplementary Note 3).

#### 4.4.5 Cell-cell communications inference

Our cell-cell communication inference method enhances biological fidelity by utilising distinct molecular modalities: transcriptomic data for ligand expression and proteomic data for receptors. This strategy directly addresses the often poor correlation between mRNA and protein levels and leverages direct cell surface protein quantification (or robust model-based predictions thereof) for a more accurate assessment of cellular receptive capacity. Specifically, ligand expression is derived from scRNA-seq. The requisite cell surface receptor proteomic data are obtained either via zero-shot prediction from scRNA-seq alone or through model fine-tuning and imputation when partial proteomic measurements (e.g., from CITE-seq) are available. Communication pathways are subsequently inferred using these modality-specific profiles and a curated consensus ligand-receptor interaction database (e.g., incorporating resources such as CellPhoneDB [62], CellChat [41], ICELLNET [63], connectome [40], and CellTalkDB [64]). A ligand (from RNA data) or a receptor (from protein data) is considered expressed within a cell type if its molecular abundance meets defined criteria (e.g., a 30% threshold percentage of expressing cells). For multimeric receptors, the coexpression of all constituent subunits, assessed at the protein level, is required for the receptor to be considered functionally present. The potential strength of interactions between cell populations is quantified based on the mean expression of participating ligands (RNA-derived) and their cognate receptors (protein-derived), accounting for the limiting subunit (protein level) in multimeric receptor complexes. Statistical significance is determined via a permutation test (e.g., 1,000 permutations), wherein cell population labels are randomly shuffled to generate an empirical null distribution for each ligand-receptor interaction score. Interactions yielding an empirical p-value below a defined significance threshold (e.g., *p* < 0.05) are deemed statistically significant. This process yields a network of putative intercellular signalling pathways robustly grounded in our modality-specific molecular assessment.

To infer intercellular signalling pathways, we first establish comprehensive singlecell transcriptomic and proteomic profiles. When only scRNA-seq data are available, our model performs zero-shot prediction of protein expression, leveraging its preestablished understanding of transcript-protein relationships. Alternatively, where paired scRNA-seq and partial proteomic data are provided, the model is fine-tuned to enhance predictions and impute expression for proteins not directly measured, thereby enriching the proteomic information for subsequent communication analysis. This strategy directly addresses the often-poor correlation between mRNA and protein levels and leverages direct cell surface protein quantification (or robust model-based predictions thereof) for a more accurate assessment of cellular receptive capacity.

Cell-cell communication is then inferred from these integrated profiles using a curated ligand-receptor interaction database, which is a consensus resource composed by multiple expert-curated ligand-receptor resources, including CellPhoneDB [62], CellChat [41], ICELLNET [63], connectome [40], and CellTalkDB [64]. A ligand or receptor is considered expressed within a cell type if its expression meets defined criteria (e.g., a minimum expression level and percentage of expressing cells). For multimeric receptors, the co-expression of all constituent subunits within the same cell population is required.

The potential strength of interactions between cell populations is quantified based on the mean expression of the participating ligands and their cognate receptors, accounting for the limiting subunit in multimeric receptor complexes. Statistical significance is assessed via a permutation test (e.g., 1,000 permutations), wherein cell population labels are randomly shuffled to generate an empirical null distribution for each ligand-receptor interaction score. An empirical p-value is subsequently derived by comparing the observed interaction score against this null distribution. Interactions falling below a defined significance threshold (e.g., *p <* 0.05) are deemed statistically significant, collectively forming a network of putative intercellular signalling pathways.

## 4.5 Code availability

The codebase for CAPTAIN is publicly available at https://github.com/iamjiboya/CAPTAIN. All analyses conducted in this paper can be reproduced using this repository at https://github.com/iamjiboya/CAPTAIN/tree/main/downstreamtasks.

4.6 Acknowledgements

We are grateful to members of the Yu laboratory and numerous colleagues for valuable comments and suggestions. This work was supported in part by the grant 2023YFF1204701 from the National Key R&D Program of China, grant 2024B1515020080 from Guangdong Basic and Applied Basic Research Foundation, grant 32470634 from the National Natural Science Foundation of China, grant KY012023362 from the Talent Research Funding Project of Guangdong Provincial People’s Hospital and grants GZNL2024A01003 and GZNL2023A02002 from the Major Project of Guangzhou National Laboratory. We acknowledge the Data Science Platform of Guangzhou National Laboratory and the Bio-medical Big Data Operating System (Bio-OS) for technical support and for providing access to the computational resources essential to this study.

## 4.7 Author contributions

F.Y. and B.J. conceived the project and designed the algorithm. B.J. implemented the algorithm. T.H. and J.W. constructed the scT&P-4M dataset. B.J., T.H., J.W., M.L., L.X., Q.Z., L.Q., Y.Z., and S.P. performed computational experiments and contributed to data interpretation. S.Z. developed the companion website. B.J., T.H., J.W., and F.Y. wrote the manuscript with input from all authors. F.Y., Y.Z., and S.P. supervised and directed the study.

