## Supplemental Files for "CAPTAIN: A multimodal foundation model pretrained on co-assayed single-cell RNA and protein"

### Supplementary Notes

#### 1 Detailed Information of Pre-trained Datasets

To establish a comprehensive benchmark for single-cell multi-omic analysis, we curated a diverse data corpus designed to rigorously test computational methods across various technological platforms and biological systems. The corpus encompasses **46 RNA + protein datasets**. This collection spans eight leading single-cell technologies, such as **CITE-seq, TEA-seq, DOGMA-seq, and Perturb-CITE-seq**, thereby ensuring a broad and representative assessment of current experimental approaches. (Supplementary Table 1).

All datasets were sourced from public repositories, with accession details provided in the "Data availability" section. To ensure data integrity and comparability, we implemented a standardized quality control pipeline for each dataset. This process leveraged the Seurat framework and strictly adhered to the filtering parameters and quality metrics, such as mitochondrial content, UMI counts, and cell viability, as described in the original publications. This approach guarantees that our analyzes are based on high-quality data that respect the specific standards of each study [1].

The curated datasets were further stratified by biological context. We provide a detailed summary of the sample distribution in various tissues and organs in Supplementary Table 2, while the breakdown by disease state, including healthy, cancer, and other conditions, is presented in Supplementary Table 3. Ultimately, this meticulously curated and annotated resource serves as a foundational benchmark for the development and validation of computational methods, enabling robust evaluation across diverse biological systems and sequencing platforms.

### 2 Standardization and Functional Annotation for Cell Surface Protein

#### Standardization Annotation for Cell Surface Protein

To enable cross-dataset integration and comparison of cell surface protein measurements across our diverse collection of single-cell RNA + protein datasets, we developed a systematic normalization pipeline to standardize cell surface protein feature names. The raw single-cell protein expression data exhibited significant heterogeneity in cell surface protein naming conventions originating from different laboratories and experimental platforms. We identified three primary sources of naming inconsistency: (1) variable prefixes reflecting different antibody cloning strategies or target epitopes (e.g., "CD45RA-BV421" vs. "CD45RA"), (2) heterogeneous suffixes containing technical information about batches, reagents, or fluorophores (e.g., "-A0251" indicating a specific batch code), and (3) irregular capitalization patterns that violated standard gene/protein nomenclature conventions (e.g., "CD3" vs. "Cd3").

To address these inconsistencies, we implemented a custom Linux-based normalization pipeline that employed regular expressions and controlled vocabulary mapping to transform raw cell surface protein labels into a standardized feature nomenclature. The pipeline systematically performed four key operations: (1) removal of all substrings following hyphens to eliminate technical details (e.g., converting "CD19-BV786" to "CD19"), (2) elimination of product codes and fluorophore tags that were irrelevant for biological interpretation, (3) standardization of all feature names to uppercase to conform with standard gene/protein nomenclature conventions, and (4) manual resolution of synonymous markers by mapping different naming variants to a unified name based on the target protein (e.g., mapping both "CD278" and "ICOS" to the standardized name "ICOS").

#### Functional Annotation for Cell Surface Protein

To create a comprehensive resource for the 382 cell surface proteins (CSPs) predictable by the CAPTAIN model, we began by integrating the protein modality from the scT&P-4M dataset. For each CSP, we systematically curated detailed annotations by leveraging a suite of established biological databases. These included Abcam's guide to human CD antigens (https://www.researchgate.net/profile/Zaffar-Equbal/publication/315478747_Baligar_et_al-2017-Hepatologysup-1/data/58d18746458515b8d285da80/Baligar-et-al-2017-Hepatologysup-1.pdf), the Human Cell Differentiation Molecules (HCDM) database for HLDA information (https://www.hcdm.org/index.php/molecule-information/Users/hutingting/Zotero/storage/WN6DY7IP/molecule-information.html), BioLegend (the source for the majority of antibodies used in high-throughput sequencing panels, https://www.biolegend.com/en-gb), the BD Biosciences CD Marker Handbook (https://www.bdbiosciences.com/content/dam/bdb/marketing-documents/cd_marker_handbook.pdf), CD-Maps (http://bioinformin.cesnet.cz/CDmaps/), NCBI (https://www.ncbi.nlm.nih.gov/), Ensembl (release 114) (https://www.ensembl.org/index.html), UniProt (https://www.uniprot.org/), GeneCards (https://www.genecards.org/), the MCE CD Antigens database (https://www.medchemexpress.com/proteins/cell-adhesion-related-cd-proteins.html?srsltid=AfmBOoq5mqT_499bcr98vR3pIbD0_D80BhKRzb_oaa9VE8egWidhpQsD), and the PanglaoDB (https://panglaodb.se/) for cell-type-specific expression data in mice.

Based on this extensive curation of functional and expression-based evidence, we systematically classified all 382 CSPs into nine key categories. This framework provides a functional map of the protein panel, encompassing core immune processes and broader physiological roles. The categories are: Cross-system Regulation, Immune Co-signaling, Cell Adhesion & Trafficking, Classical Signaling Receptors, Antigen Presentation, Cell Lineage & Identity Markers, Tumor Metabolism & Proliferation, and Immune Recognition & Development. A final category, Others, includes proteins with currently unclassified or miscellaneous functions.

For each of the 382 CSPs, we compiled their full name, gene name, aliases, primary molecular function (receptor, ligand, etc.), and associations with key immune regulators like MHC and interleukins. To elucidate their biological context, we aggregated expression profiles across a wide range of cell types (e.g., T cells, B cells, macrophages, epithelial cells, tumor cells). We documented their relevance to various tissues, diseases, and core biological processes such as immune recognition, cell activation, and immune checkpoint regulation.

For detailed metrics on the normalization process, including the number of features processed, mapping success rates, and resolution of synonymous markers, see Supplementary Tables 4 - 8. The standardized feature names established through this pipeline have been used consistently throughout our analyses to ensure accurate cross-dataset comparisons of cell surface protein expression patterns.

#### 3 Evaluation Metric Calculations

##### 3.1 Cell Type Assignment

We used the standard classification metrics *Accuracy*, *Precision*, *Recall*, and $MacroF1$ to evaluate cell type assignment performance. The *Accuracy*, *Precision*, *Recall*, and $MacroF1$ scores are calculated from true positives ( $tp$ ), false positives ( $fp$ ), and false negatives ( $fn$ ) globally or averaged per class $c$ of $N_c$ cells.

The *Accuracy*, *Precision* and *Recall* scores are calculated as follows:

$$Accuracy = \frac{\sum_{c\in C} tp_c}{\sum_{c\in C} N_c}, \quad Precision = \frac{1}{|C|}\sum_{c\in C}\frac{tp_c}{tp_c + fp_c}, \quad Recall = \frac{1}{|C|}\sum_{c\in C}\frac{tp_c}{tp_c + fn_c}.$$

The $Macro - F1$ score is calculated per cell type $c$ first and averaged across cell types:

$$Macro - F1 = \frac{1}{|C|}\sum_{c\in C} F1_c, \quad \text{where} \quad F1_c = \frac{2 \times Precision_c \times Recall_c}{Precision_c + Recall_c}$$

The above metrics are calculated using scikit-learn's implementations.

##### 3.2 Single-cell integration

We adopted the evaluation metric calculations outlined by Luecken et al [2]. in their benchmark study. Each metric is described below.

**Normalized Mutual Information:** To quantify the concurrence between the cell type labels based on ground truth and the Louvain cluster labels obtained from integrated cell embeddings, we computed the normalized mutual information (NMI) score. The Louvain clustering was conducted across resolutions ranging from 0.1 to 2, with increments of 0.1. The best score will be selected. The NMI score for cell types, referred to as $\boldsymbol{NMI}_{cell}$, ranges between 0 and 1 , where a higher score indicates a better match of cell types.

**Adjusted Rand Index:** The adjusted rand index (ARI) was employed to assess both the agreement between the annotated labels and the MNI-optimized Louvain clusters. Furthermore, the rand index was adjusted to account for randomly correct labels. The ARI score for cell types, denoted as $\boldsymbol{ARI}_{cell}$, ranges from 0 to 1 , where 0 corresponds to random labeling and 1 represents a perfect match.

**Average Silhouette Width:** The silhouette width assesses the relationship between a cell's within-cluster distances and its distances to the closest cluster boundaries. By averaging the silhouette widths of all cells, we calculate the average silhouette width (ASW) score. This score ranges from -1 to 1 , where a score of 1 indicates well-separated clusters, while scores from -1 to 0 suggest overlapping clusters and misclassification.

For evaluating cell type clustering, we compute the ASW score based on cell type labels, represented as $\boldsymbol{ASW}_{cell}$. To obtain this score, we utilize the following formula:

$$ASW_{cell} = (ASW_C + 1)/2$$

Here, $C$ represents the cell types.

Regarding batch mixing evaluation, we calculate the ASW score considering batch labels and adjust it by subtracting 1. This score is denoted as $\boldsymbol{ASW}_{batch}$. The calculation is as follows:

$$ASW_{\text{batch}} = 1 - |ASW_B|$$

Both $\boldsymbol{ASW}_{cell}$ and $\boldsymbol{ASW}_{batch}$ have values between 0 and 1. Higher scores indicate better cell-type clustering or batch-mixing performance.

**Graph Connectivity:** The graph connectivity metric quantifies the average proportion of cells within each cell type that are connected through a kNN (k-nearest neighbors) graph. For every cell identity $c$ in the set $C$, we compute the size of the largest connected component using kNN among cells exclusively belonging to identity $c$. This value is divided by the total number of cells with identity $c$ to obtain a normalized measure. The *GraphConn* score is then reported as the average across all cell types:

$$GraphConn = \frac{1}{|C|} \sum_{c \in C} \frac{\left| LCC\left(G_c^{kNN}\right) \right|}{N_c}$$

Here, $LCC$ represents the largest connected component, and $N$ denotes the number of cells of each celltype.

**Aggregated Metrics:** The aggregated metric $\boldsymbol{AvgBIO}$ calculates the average of biological conservation metrics:

$$AvgBIO = (ARI_{cell} + NMI_{cell} + ASW_{cell})/3$$

Similarly, the aggregated metric $\boldsymbol{AvgBATCH}$ computes the average of batch mixing metrics:

$$AvgBATCH = (ASW_{batch} + GraphConn)/2$$

In accordance with the convention established in [2], an Overall metric is derived as the weighted average of 0.6 × AvgBIO + 0.4 × AvgBATCH.

#### 4 Comparison to Existing Single-Cell Foundation Models

The advent of Transformer architectures has spurred the development of foundation models for single-cell biology, aiming to create versatile tools for a wide array of analytical tasks. This section compares CAPTAIN with several state-of-the-art models, including **scFoundation** [**3**], **scBERT** [**4**], **scGPT**, and **Geneformer** [**5**], across various dimensions from pre-training data to downstream applications (Supplementary Table 14).

**Pre-training Data and Scale.** The scale and diversity of pre-training data are fundamental to a model's generalizability. Among the compared models, scFoundation is pre-trained on the largest dataset, comprising 50 million cells from an extensive collection of public data (HCA, GEO, hECA, etc.) spanning 264 tissues. Geneformer follows with 30 million cells across 39 organs. In comparison, CAPTAIN utilizes a dataset of 4 million cells from 26 tissues, while scGPT and scBERT are trained on 10 million and 1 million cells, respectively. Notably, while most models focus solely on human data, CAPTAIN and Geneformer are pre-trained on data from both Human and Mouse, broadening their cross-species applicability.

**Model Architecture and Pre-training Strategy.** The models exhibit significant differences in their architectural design. CAPTAIN employs a unique Encoder (cross-model) architecture, whereas scFoundation utilizes an Asymmetric Encoder-Decoder structure. In contrast, scBERT, scGPT, and Geneformer adopt a more conventional Encoder Only design. Model size also varies, with scFoundation having the largest parameter count at ~ 100M, followed by CAPTAIN (~ 60M), scGPT (~ 50M), Geneformer (~ 30M), and scBERT (~ 10M). Regarding input data, CAPTAIN, scBERT, and scGPT process binned normalized expression values. scFoundation, however, uses continuous normalized expression values, and Geneformer takes ranked normalized expression values. A key distinction for CAPTAIN is its input of not only ~ 3,000 non-zero genes but also 382 cell surface protein features, enabling native multi-omic analysis from the input level. This is reflected in its pre-training tasks, which, in addition to the common Masked gene value prediction, include specialized objectives like Protein expression prediction and Protein expression interval prediction. This contrasts with other models that primarily focus on masked value prediction, although scFoundation incorporates a read-depth aware mechanism.

**Downstream Task Capabilities.** The ultimate utility of a foundation model lies in its performance across diverse downstream tasks. All compared models demonstrate proficiency in Cell type annotation. However, CAPTAIN showcases a unique and strong capability in protein-centric and multi-omic analyses. It is the only model among the group explicitly shown to perform Protein prediction, Protein imputation, and Cellular communication analysis. Furthermore, similar to scGPT, CAPTAIN supports Multi-omic integration, a task not addressed by scFoundation, scBERT, or Geneformer.

Conversely, some models have their own specialized strengths. For instance, scFoundation excels at Read-depth enhancement and Drug response prediction. Geneformer is uniquely capable of Cell fate prediction. While CAPTAIN is generic, it does not currently support perturbation prediction tasks solved by scFoundation, scGPT, and Geneformer. We will progress to add downstream tasks to achieve more fine-tuning projects. This comparative analysis highlights CAPTAIN's distinct position as a powerful multi-modal foundation model with a specialized focus on integrating transcriptomic and proteomic data.

### Supplementary Tables

Supplementary Table 1: Detailed Information of Pre-trained Datasets (Given its extensive size, the complete table is provided as Table 1 in the Supplementary File.)

| **Tissue** | **Sample** | **Cells** |
|---|---|---|
| **Blood** | **86** | **2,250,157** |
| Bone Marrow | 39 | 551,073 |
| Lung | 41 | 396,267 |
| Lymphoid | 43 | 254,010 |
| Skin | 3 | 243,566 |
| Brain | 4 | 211,059 |
| Intestine | 1 | 150,363 |
| Liver | 10 | 93,578 |
| Others | 4 | 37,915 |
| Kidney | 6 | 33,483 |
| Bone | 1 | 12,226 |
| Breast | 6 | 11,141 |
| Pancreas | 2 | 8,330 |
| Spleen | 3 | 7,377 |
| Total | 249 | 4,260,545 |

Supplementary Table 2: A summary of the pretrained dataset. The following table provides a detailed breakdown of the pre-training data cohort used in this study. The number of unique samples and the total cell count for each constituent tissue type are listed. In total, the dataset includes 4,531,996 cells from 258 samples. The cohort is predominantly composed of cells from Peripheral Blood Mononuclear Cells (PBMCs).

| Disease | Sample | Cells |
|---|---|---|
| Healthy | 130 | 1,397,193 |
| **Coronavirus Disease 2019** | **38** | **1,234,362** |
| Cancer | 38 | 1,035,790 |
| Acute Respiratory Distress Syndrome | 24 | 193,285 |
| Ulcerative Colitis | 1 | 150,363 |
| Epilepsy | 1 | 84,821 |
| Hepatitis B Virus-Associated Chronic Liver Disease | 6 | 55,614 |
| Immunological Disease | 6 | 37,589 |
| Hypertension | 2 | 30,073 |
| Psoriasis | 2 | 28,704 |
| Coronary Artery Disease | 1 | 12,751 |
| Total | 249 | 4,260,545 |

Supplementary Table 3: Clinical status classification of samples in the study cohort. For disease-specific analysis, all samples were rigorously categorized into 11 distinct groups based on their clinical diagnosis of origin, comprising a healthy control group and ten major disease types. These labels highlight the breadth and heterogeneity of our training dataset.

Supplementary Table 4: Cell Surface Protein vocabulary (Given its extensive size, the complete table is provided as Table 4 in the Supplementary File.)

Supplementary Table 5: Comparison of the original names of cell surface proteins with their names in our dictionary (Given its extensive size, the complete table is provided as Table 5 in the Supplementary File.)

Supplementary Table 6: Feature annotations of Cell Surface Protein in our dictionary (Given its extensive size, the complete table is provided as Table 6 in the Supplementary File.)

Supplementary Table 7: Function categories of each Cell Surface Protein in our dictionary (Given its extensive size, the complete table is provided as Table 7 in the Supplementary File.)

Supplementary Table 8: Function summary of Cell Surface Protein in our dictionary (Given its extensive size, the complete table is provided as Table 8 in the Supplementary File.)

#### Benchmarking Results on Downstream Tasks

**Pearson correlation and RMSE results for MALT dataset from 10x Genomics**

| | CAPTAIN | | CAPTAIN_zero-shot | | Seurat [6] | | sciPENN [7] | | TotalVI [8] | |
|---|---|---|---|---|---|---|---|---|---|---|
| CSP_name | Pearson | RMSE | Pearson | RMSE | Pearson | RMSE | Pearson | RMSE | Pearson | RMSE |
| CD8a | 0.823 | 0.575 | 0.691 | 0.888 | −0.028 | 1057.712 | 0.456 | 3.106 | 0.658 | 0.827 |
| CD14 | 0.803 | 0.629 | 0.678 | 1.090 | 0.051 | 198.195 | 0.293 | 2.583 | −0.043 | 1.444 |
| CD15 | 0.254 | 0.972 | 0.050 | 1.146 | −0.013 | 270.206 | −0.195 | 2.808 | 0.571 | 0.927 |
| CD16 | 0.843 | 0.559 | 0.577 | 3.059 | −0.003 | 216.923 | −0.267 | 2.856 | 0.292 | 1.190 |
| CD19 | 0.899 | 0.441 | 0.691 | 1.205 | 0.371 | 1316.006 | 0.837 | 5.566 | −0.110 | 1.490 |
| CD25 | 0.424 | 0.910 | 0.420 | 1.024 | 0.007 | 436.964 | 0.269 | 3.383 | 0.626 | 0.865 |
| CD45RA | 0.793 | 0.617 | 0.693 | 0.867 | 0.018 | 530.745 | 0.613 | 4.957 | 0.305 | 1.179 |
| CD45RO | 0.614 | 0.805 | 0.396 | 0.942 | −0.082 | 1058.472 | 0.670 | 4.180 | 0.570 | 0.928 |
| TIGIT | 0.402 | 0.913 | 0.384 | 0.930 | −0.019 | 340.734 | 0.178 | 3.278 | 0.112 | 1.333 |
| CD127 | 0.796 | 0.610 | 0.714 | 0.792 | −0.148 | 214.657 | 0.245 | 3.040 | 0.087 | 1.352 |

**Pearson correlation and RMSE results for MNC [7] dataset**

| | CAPTAIN | | CAPTAIN_zero-shot | | Seurat | | sciPENN | | TotalVI | |
|---|---|---|---|---|---|---|---|---|---|---|
| CSP_name | Pearson | RMSE | Pearson | RMSE | Pearson | RMSE | Pearson | RMSE | Pearson | RMSE |
| CD3 | 0.934 | 0.132 | 0.796 | 2.851 | 0.900 | 0.348 | 0.790 | 4.299 | 0.802 | 0.629 |
| CD4 | 0.917 | 0.162 | 0.803 | 1.626 | 0.849 | 0.462 | 0.677 | 4.271 | 0.707 | 0.765 |
| CD8 | 0.806 | 0.373 | 0.620 | 1.293 | 0.663 | 0.820 | 0.494 | 4.061 | 0.510 | 0.989 |
| CD2 | 0.856 | 0.281 | 0.733 | 2.880 | 0.840 | 0.429 | 0.675 | 4.853 | 0.657 | 0.828 |
| CD45RA | 0.869 | 0.253 | 0.742 | 1.432 | 0.847 | 0.701 | 0.525 | 6.157 | 0.502 | 0.998 |
| CD57 | 0.725 | 0.516 | 0.331 | 0.972 | 0.634 | 0.302 | 0.657 | 3.893 | 0.368 | 1.124 |
| CD16 | 0.767 | 0.439 | 0.155 | 1.527 | 0.679 | 0.101 | 0.677 | 2.592 | 0.370 | 1.122 |
| CD14 | 0.905 | 0.183 | 0.810 | 0.894 | 0.892 | 0.147 | 0.775 | 2.811 | 0.802 | 0.630 |
| CD11c | 0.914 | 0.166 | 0.859 | 1.213 | 0.928 | 0.176 | 0.854 | 2.712 | 0.834 | 0.575 |
| CD19 | 0.892 | 0.213 | 0.784 | 0.742 | 0.962 | 0.194 | 0.809 | 2.883 | 0.720 | 0.748 |

Supplementary Table 9: Performance benchmark of CAPTAIN for single-cell protein imputation and expansion. A comparative analysis of our method, CAPTAIN, against several state-of-the-art approaches: Seurat [6], sciPENN [7] and TotalVI [8]. A zero-shot variant of our method (CAPTAIN_zero-shot), which performs prediction without access to protein information in the reference data. The evaluation was conducted across four public CITE-seq datasets: Human MALT from 10x Genomics, Human MNC [7], Mouse PBMC [9] and Human PBMC [10]. Performance was quantified by calculating the Pearson correlation coefficient (Pearson) and the Root Mean Square Error (RMSE) between the imputed and ground-truth protein abundance values for every protein in the test sets. For more information on the other datasets, please see Supplementary Table 9 in our Supplementary file.

| Modality | Dataset | Model | Classification Metrics | | | |
|---|---|---|---|---|---|---|
| | | | *Accuracy* | *Precision* | *Recall* | *MacroF1* |
| scRNA | 10x Multiome PBMC | CAPTAIN (fine-tuned) | 0.962 | 0.961 | **0.937** | **0.947** |
| | | scGPT( fine-tuned) [11] | 0.945 | 0.867 | 0.876 | 0.868 |
| | | Seurat | **0.972** | **0.970** | 0.923 | 0.943 |
| | Multi-tissue Immune Cells [12] | CAPTAIN (fine-tuned) | **0.922** | **0.898** | **0.920** | **0.905** |
| | | scGPT( fine-tuned) | 0.915 | 0.875 | 0.891 | 0.877 |
| | | Seurat | 0.790 | 0.848 | 0.732 | 0.761 |
| | Integrated COVID-19 (H2C) [7, 13, 14] | CAPTAIN (fine-tuned) | **0.856** | **0.847** | **0.807** | **0.809** |
| | | scGPT( fine-tuned) | 0.821 | 0.822 | 0.791 | 0.788 |
| | | Seurat | 0.799 | 0.784 | 0.732 | 0.733 |
| | Integrated COVID-19 (C2C) [7, 13, 14] | CAPTAIN (fine-tuned) | **0.881** | **0.847** | **0.818** | **0.830** |
| | | scGPT( fine-tuned) | 0.856 | 0.807 | 0.793 | 0.794 |
| | | Seurat | 0.827 | 0.791 | 0.716 | 0.741 |
| | Myasthenia Gravis Thymoma/Thymus [15] | CAPTAIN (fine-tuned) | **0.871** | **0.807** | **0.791** | **0.786** |
| | | scGPT( fine-tuned) | 0.836 | 0.714 | 0.644 | 0.639 |
| | | Seurat | 0.848 | 0.793 | 0.684 | 0.707 |
| CITE-seq | Human PBMC [10] | CAPTAIN (fine-tuned) | **0.970** | **0.954** | **0.960** | **0.957** |
| | | scGPT( fine-tuned) | 0.719 | 0.479 | 0.502 | 0.467 |
| | | Seurat | 0.471 | 0.486 | 0.336 | 0.499 |
| | COVID-19 Multi-modal PBMC [16] | CAPTAIN (fine-tuned) | **0.897** | **0.820** | **0.755** | **0.769** |
| | | scGPT( fine-tuned) | 0.701 | 0.293 | 0.282 | 0.276 |
| | | Seurat | 0.670 | 0.773 | 0.273 | 0.668 |
| | Integrated COVID-19 [7, 13, 14] | CAPTAIN (fine-tuned) | **0.871** | **0.829** | **0.823** | **0.821** |
| | | scGPT( fine-tuned) | 0.171 | 0.012 | 0.071 | 0.021 |
| | | Seurat | 0.043 | 0.021 | 0.025 | 0.083 |
| | BMMC [17] | CAPTAIN (fine-tuned) | **0.828** | **0.708** | **0.745** | **0.718** |
| | | scGPT( fine-tuned) | 0.803 | 0.148 | 0.202 | 0.161 |
| | | Seurat | 0.778 | 0.310 | 0.119 | 0.545 |
| | BMMC Subset | CAPTAIN (fine-tuned) | **0.836** | **0.730** | **0.745** | **0.730** |
| | | scGPT( fine-tuned) | 0.434 | 0.033 | 0.077 | 0.047 |
| | | Seurat | 0.521 | 0.186 | 0.112 | 0.610 |

Supplementary Table 10: Benchmark Results for Cell Type Annotation. The performance of CAPTAIN for cell type annotation, benchmarked against scGPT [11] and Seurat [**?**]. The evaluation was conducted on a diverse collection of eight datasets, grouped by data modality. The scRNA-seq datasets include: 10x Multiome PBMC from 10x Genomics, Multi-tissue Immune Cells [12], Integrated COVID-19 (H2C) [7, 13, 14], Integrated COVID-19 (C2C) [7, 13, 14] and Myasthenia Gravis Thymoma/Thymus [15]. The multi-modal (CITE-seq) datasets include: Human PBMC [10], a COVID-19 Multi-modal PBMC [16], Integrated COVID-19 [7, 13, 14] and BMMC [17] which combines the subset datasets of BMMC with $CD4^+$ T cell and $CD8^+$ T cell. Performance was assessed using four standard classification metrics: Accuracy, Precision, Recall, and Macro-F1 score.

| Dataset | Model | Biological Conservation | | | | Batch Correction | | | |
|---|---|---|---|---|---|---|---|---|---|
| | | **AvgBIO** | $NMI_{cell}$ | $ARI_{cell}$ | $ASW_{cell}$ | **AvgBATCH** | $ASW_{batch}$ | *GraphConn* | **Overall** |
| PBMCs Healthy | CAPTAIN (fine-tuned) | **0.832** | 0.923 | 0.937 | 0.636 | **0.953** | 0.912 | 0.994 | **0.880** |
| | scGPT (fine-tuned) | 0.725 | 0.785 | 0.735 | 0.656 | **0.944** | 0.899 | 0.988 | 0.813 |
| | Seurat | 0.569 | 0.735 | 0.608 | 0.363 | 0.718 | 0.534 | 0.902 | 0.628 |
| | Harmony [18] | 0.649 | 0.852 | 0.805 | 0.290 | 0.717 | 0.540 | 0.895 | 0.676 |
| PBMCs SARS-COV-2 | CAPTAIN (fine-tuned) | **0.655** | 0.733 | 0.694 | 0.537 | **0.839** | 0.857 | 0.822 | **0.729** |
| | scGPT (fine-tuned) | 0.523 | 0.608 | 0.464 | 0.496 | 0.857 | 0.920 | 0.795 | 0.657 |
| | Seurat | 0.384 | 0.607 | 0.516 | 0.029 | 0.631 | 0.498 | 0.764 | 0.483 |
| | Harmony | 0.379 | 0.604 | 0.507 | 0.026 | 0.631 | 0.497 | 0.764 | 0.480 |
| BMMCs Healthy | CAPTAIN (fine-tuned) | **0.664** | 0.700 | 0.675 | 0.616 | **0.858** | 0.807 | 0.910 | **0.742** |
| | scGPT (fine-tuned) | 0.592 | 0.646 | 0.488 | 0.642 | 0.868 | 0.799 | 0.938 | 0.703 |
| | Seurat | 0.354 | 0.530 | 0.529 | 0.005 | 0.591 | 0.510 | 0.672 | 0.449 |
| | Harmony | 0.337 | 0.470 | 0.555 | −0.014 | 0.576 | 0.503 | 0.648 | 0.433 |

Supplementary Table 11: scRNA-seq Integration Benchmark Results. CAPTAIN was benchmarked with scGPT, Seurat, and Harmony on the PBMCs Healthy (8 batches), PBMCs SARS-COV-2 (20 batches) and BMMCs Healthy (13 batches) datasets for cell type clustering and batch correction performance. The CAPTAIN (fine-tuned) is a smaller four-layer transformer model trained from random parameter initialization. We present three aggregate scores: $AvgBIO$, $AvgBATCH$ and $Overall$. These aggregate scores were calculated from three detailed biological conservation metrics ($NMI_{cell}$, $ARI_{cell}$, $ASW_{cell}$) and two batch correction metrics ($ASW_{batch}$, $GraphConn$).

| Dataset | Model | Biological Conservation | | | | Batch Correction | | | |
|---|---|---|---|---|---|---|---|---|---|
| | | **AvgBIO** | $NMI_{cell}$ | $ARI_{cell}$ | $ASW_{cell}$ | **AvgBATCH** | $ASW_{batch}$ | *GraphConn* | **Overall** |
| Integrated COVID-19 | CAPTAIN (fine-tuned) | **0.657** | 0.742 | 0.691 | 0.538 | - | - | - | - |
| | scGPT (fine-tuned) | 0.626 | 0.710 | 0.608 | 0.545 | - | - | - | - |
| | TotalVI [8] | 0.607 | 0.689 | 0.608 | 0.522 | - | - | - | - |
| | Scanpy [19] | 0.519 | 0.589 | 0.475 | 0.494 | - | - | - | - |
| TEA-seq + ECCITE-seq + CITE-seq [10] | CAPTAIN (fine-tuned) | **0.567** | 0.638 | 0.408 | 0.657 | **0.968** | 0.943 | 0.992 | **0.728** |
| | scGPT (fine-tuned) | 0.517 | 0.548 | 0.339 | 0.664 | 0.951 | 0.913 | 0.988 | 0.690 |

Supplementary Table 12: scMultiomic Integration Benchmark Results. For the paired Human PBMCs Covid dataset, CAPTAIN was benchmarked with scGPT [11], TotalVI [8] and Scanpy [19] for cell type clustering performance evaluated on four biological conservation metrics. For the multi-omics and multi-batch TEA-seq + ECCITE-seq + CITE-seq [10] dataset, CAPTAIN was benchmarked with scGPT (fine-tuned) respectively on cell type clustering and multi-omic integration performance.

| Ligand | Receptor | CAPTAIN | NATMI | Connectome | CellChat | CellphoneDB | SingleCellSingleR | PMID |
|---|---|---|---|---|---|---|---|---|
| CD48 | CD244 | 1 | 0 | 0 | 0 | 0 | 0 | 15210772; 27581174; 16585556 |
| YBX1 | EGFR | 1 | 0 | 0 | 0 | 0 | 0 | 19483673; |
| CD40LG | CD47 | 1 | 0 | 0 | 0 | 0 | 0 | 36768931 |
| AREG | MERTK | 1 | 0 | 0 | 0 | 0 | 0 | no |
| VEGFB | MERTK | 1 | 0 | 0 | 0 | 0 | 0 | 39145452; |
| MIF | CD74 | 1 | 0 | 0 | 0 | 0 | 0 | 18490733; |
| FLT3LG | MERTK | 1 | 0 | 0 | 0 | 0 | 0 | 11159533; 26851220 |
| CD99 | CD81 | 1 | 0 | 0 | 0 | 0 | 0 | 23858057; 29305692 |
| COPA | CD74 | 1 | 0 | 0 | 0 | 0 | 0 | 25894502 |
| CMTM8 | EGFR | 1 | 0 | 0 | 0 | 0 | 0 | 32268840 |
| MIF | EGFR | 1 | 0 | 0 | 0 | 0 | 0 | 39233830 |
| AREG | EGFR | 1 | 0 | 0 | 0 | 0 | 1 | 25692699; 30042827 |
| VEGFB | EGFR | 1 | 0 | 0 | 0 | 0 | 2 | 21439312; 31717527 |
| ARF4 | EGFR | 1 | 0 | 0 | 0 | 0 | 3 | no |
| TGFB1 | EGFR | 1 | 0 | 0 | 0 | 0 | 4 | 20934403; 29215776 |
| FLT3LG | EGFR | 1 | 0 | 0 | 0 | 0 | 5 | 7535149 |
| PTPN6 | EGFR | 1 | 0 | 0 | 0 | 0 | 6 | 35186080 |
| CD40LG | CD83 | 1 | 0 | 0 | 0 | 0 | 7 | 17301951; 9182676 |
| ANXA1 | EGFR | 1 | 0 | 0 | 0 | 0 | 8 | 23267026; 28485170 |
| CLEC2D | CD161 | 1 | 0 | 0 | 0 | 0 | 9 | 16339512; 18453569 |
| TGFB1 | SDC1 | 1 | 0 | 0 | 0 | 0 | 10 | no |
| CD40LG | CD80 | 1 | 0 | 0 | 0 | 1 | 11 | no |
| HLA-A | CD80 | 0 | 0 | 0 | 0 | 0 | 12 | no |
| HLA-A | CD28 | 0 | 0 | 0 | 0 | 0 | 13 | no |
| CD40LG | CD80 | 0 | 0 | 0 | 0 | 0 | 14 | no |
| TGFB1 | CD28 | 0 | 0 | 0 | 0 | 0 | 15 | 32228232 |
| CD40LG | CD4 | 0 | 0 | 0 | 0 | 0 | 16 | 29476811 |

Supplementary Table 13: Literature curated of the inferred intercellular communications by CAPTAIN, NATMI, Connectome, CellChat, CellphoneDB and SingleCellSingleR between $CD4^+$ Naive T and Natural Killer (NK) cells.

Supplementary Table 14: Comparison to existing single-cell foundation models (Given its extensive size, the complete table is provided as Table 14 in the Supplementary File.)

### Supplementary Figures

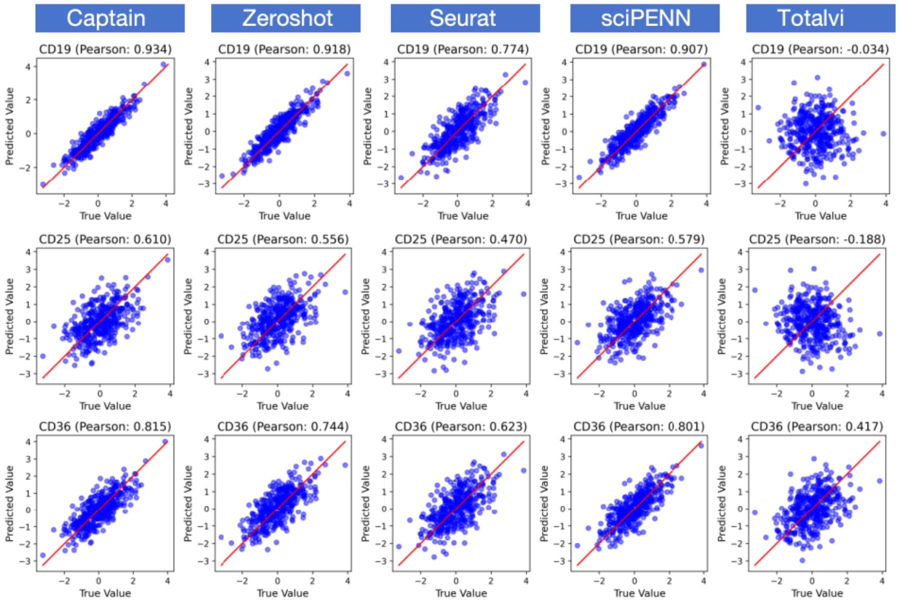


Supplementary Figure 1: Comparison of predicted vs. true protein expression across different methods (columns). The Pearson correlation for each cell surface protein is shown in the plot title. For all three proteins, CAPTAIN yields the highest correlation, outperforming other benchmarked methods.

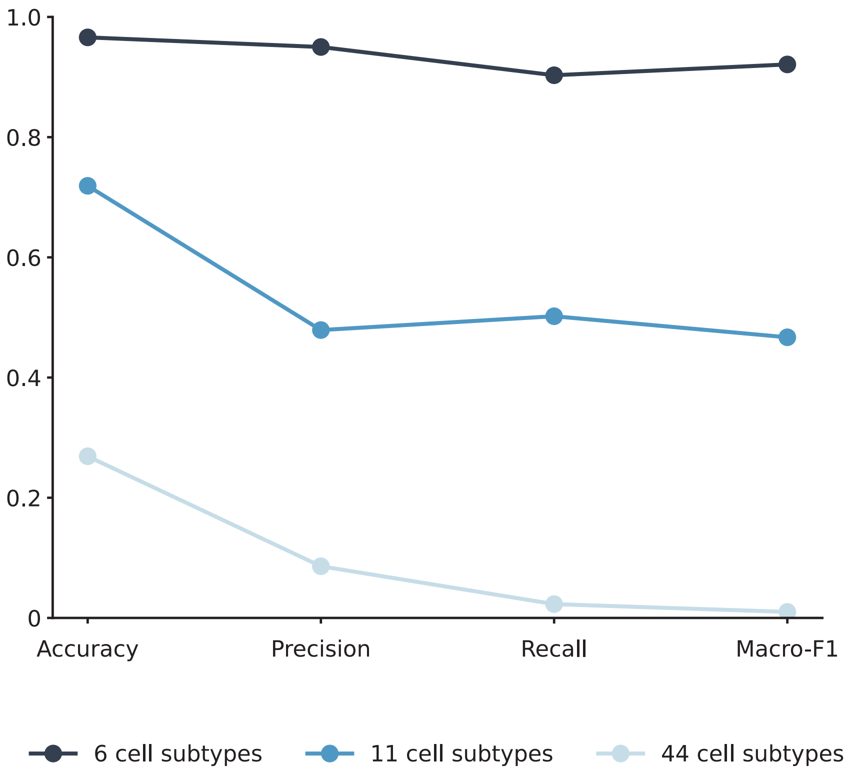


Supplementary Figure 2: Performance of scGPT for cell type annotation of multi-omics datasets declines with increasing granularity of cell subtypes. Quantitative performance metrics (Accuracy, Precision, Recall, and Macro-F1 score) for scGPT's cell type annotation on datasets with distinct cell subtypes.

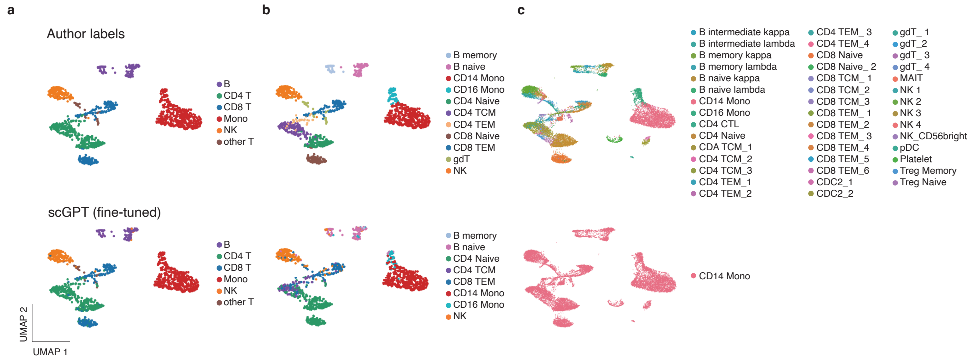


Supplementary Figure 3: UMAP Visualization Comparing scGPT Predictions with Ground Truth: Model Performance Degrades as the Number of Cell Subtypes Increases. **(a)** UMAP visualizations comparing ground truth cell types (left) and scGPT's predictions (right) for a dataset with 6 major cell subtypes. At this level of granularity, the model's predictions show high concordance with the ground truth labels. **(b)** UMAP visualizations for a dataset with 11 cell subtypes. The left panel shows the ground truth labels, while the right panel shows scGPT's predictions. The model begins to show reduced performance, failing to identify certain subtypes (e.g., gdT, CD8 Naive) that are present in the ground truth. **(c)** UMAP visualizations for a highly granular dataset with 44 distinct cell subtypes. The left panel displays the diverse ground truth cell populations. The right panel demonstrates a significant degradation in scGPT's performance, where the model largely fails to resolve the subtypes and misclassifies the majority of cells as a single type (CD14 Mono).

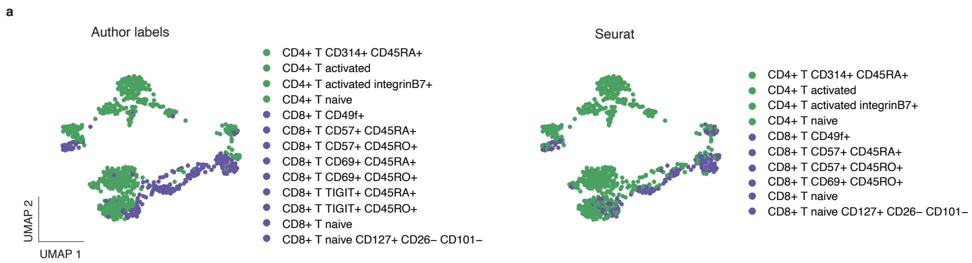


Supplementary Figure 4: A UMAP comparison of ground truth annotations (left) versus predictions by Seurat (right) on a dataset of CD4+ and CD8+ T cell subtypes. The results show that the Seurat method fails to identify several of the finer CD8+ T cell subtypes.

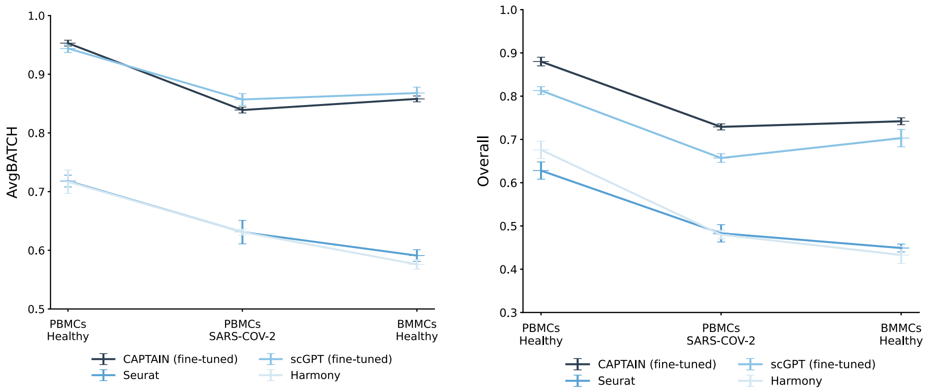


Supplementary Figure 5: Quantitative benchmark of CAPTAIN's batch correction performance against other methods. Line plots comparing the performance of four integration methods (CAPTAIN (fine-tuned), scGPT (fine-tuned), Seurat, and Harmony) across three distinct datasets: healthy PBMCs, SARS-CoV-2 PBMCs, and healthy BMMCs. The AvgBATCH score (left) quantifies the effectiveness of batch effect removal, while the Overall score (right) is a composite metric evaluating both batch correction and the preservation of biological variance (AvgBIO). A higher score indicates better performance in both metrics. The results demonstrate that CAPTAIN consistently achieves the highest Overall score across all datasets, indicating its superior ability to remove batch effects while retaining crucial biological information compared to the other benchmarked methods.

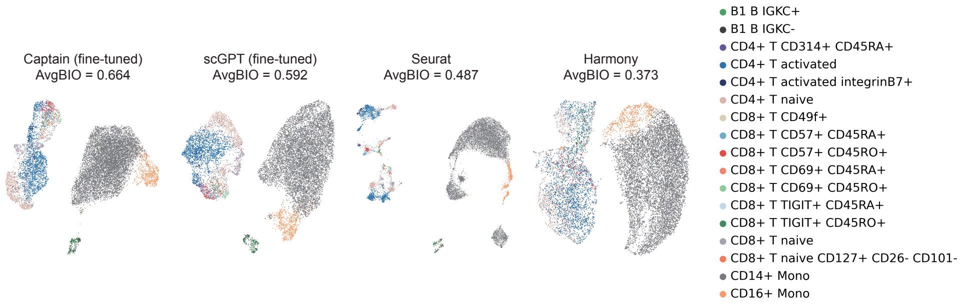


Supplementary Figure 6: UMAP visualizations comparing the performance of four different integration methods (CAPTAIN (fine-tuned), scGPT (fine-tuned), Seurat, and Harmony) on a paired gene expression and protein abundance dataset from bone marrow mononuclear cells (BMMCs).

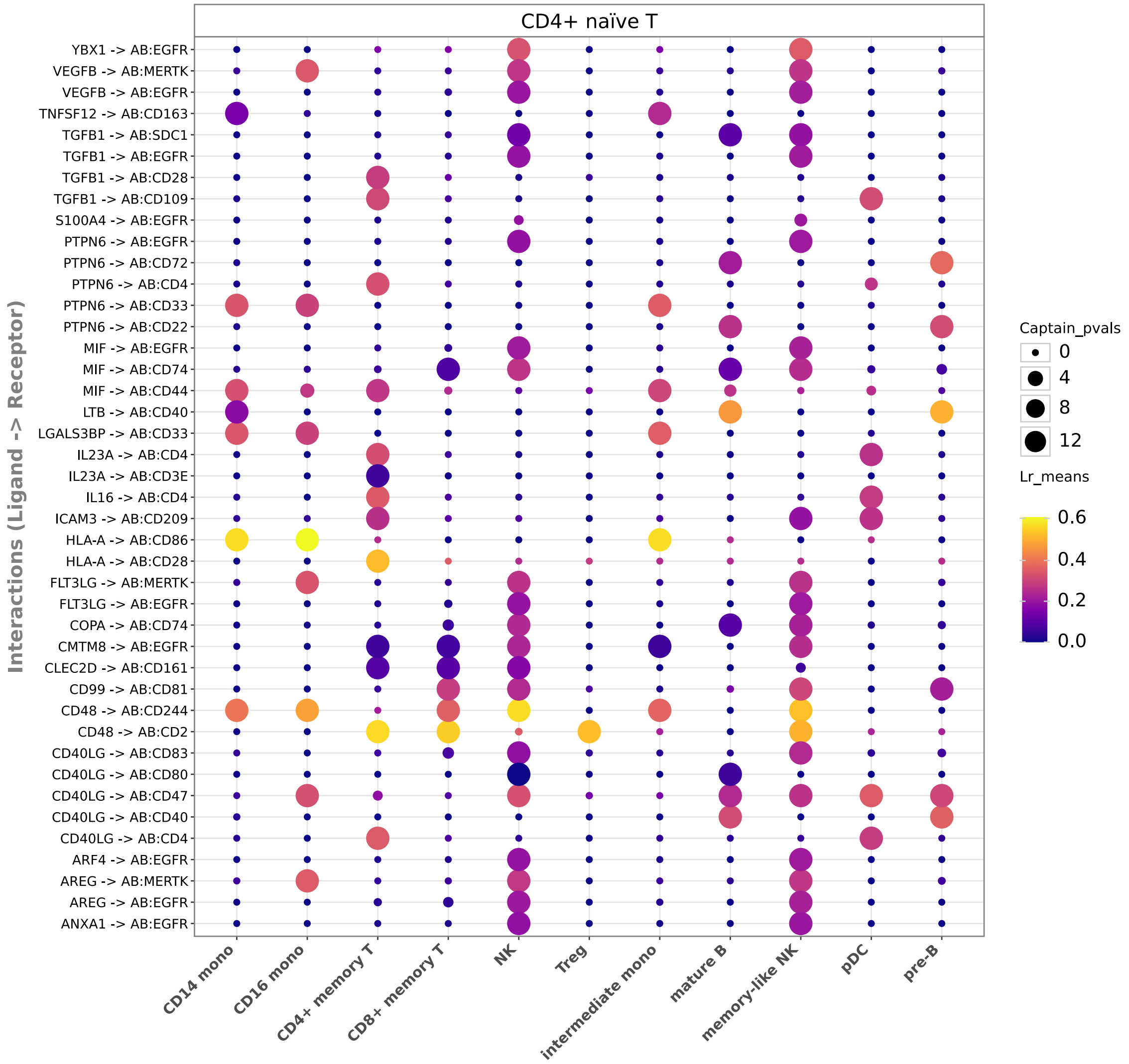


Supplementary Figure 7: A dot plot summarizing the significant intercellular communications originating from CD4+ Naïve T cells (source) to various other immune cell populations (targets), highlighting strong and significant interactions between CD4+ Naïve T cells and Natural Killer (NK) cells.

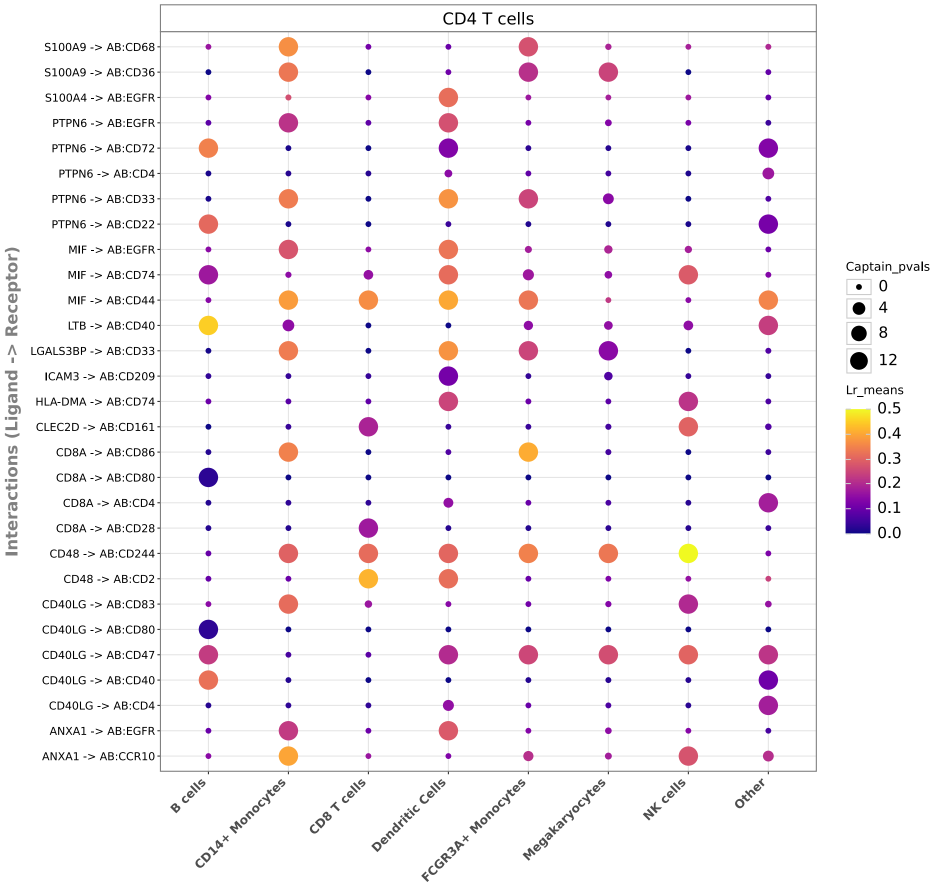


Supplementary Figure 8: A dot plot summarizing the significant intercellular communications originating from CD4 T cells (source) to various other immune cell populations (targets), highlighting strong and significant interactions between CD4 T cells and Dendritic Cellsr (DC) cells.

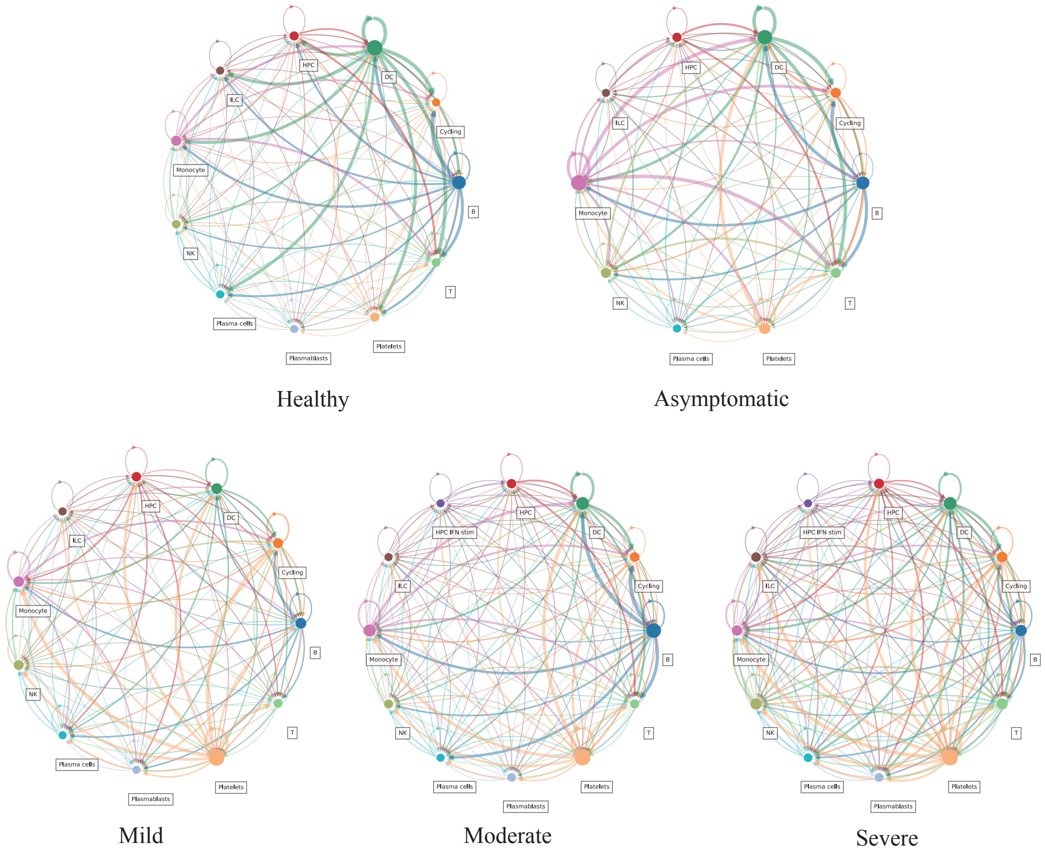


Supplementary Figure 9: Dynamics of intercellular communication networks across different COVID-19 disease states. Circle plots display the cell communication networks inferred by CAPTAIN for states ranging from Healthy to Severe COVID-19. The thickness of the lines connecting cell types corresponds to the overall interaction strength. As disease severity increases, communication between Platelets and Monocytes becomes progressively stronger, aligning with the known importance of this interaction in severe COVID-19.
